# Stochastic neural field model of stimulus-dependent variability in cortical neurons

**DOI:** 10.1101/514315

**Authors:** Paul C. Bressloff

## Abstract

We use stochastic neural field theory to analyze the stimulus-dependent tuning of neural variability in ring attractor networks. We apply perturbation methods to show how the neural field equations can be reduced to a pair of stochastic nonlinear phase equations describing the stochastic wandering of spontaneously formed tuning curves or bump solutions. These equations are analyzed using a modified version of the bivariate von Mises distribution, which is well-known in the theory of circular statistics. We first consider a single ring network and derive a simple mathematical expression that accounts for the experimentally observed bimodal (or M-shaped) tuning of neural variability. We then explore the effects of inter-network coupling on stimulus-dependent variability in a pair of ring networks. These could represent populations of cells in two different layers of a cortical hypercolumn linked via vertical synaptic connections, or two different cortical hypercolumns linked by horizontal patchy connections within the same layer. We find that neural variability can be suppressed or facilitated, depending on whether the inter-network coupling is excitatory or inhibitory, and on the relative strengths and biases of the external stimuli to the two networks. These results are consistent with the general observation that increasing the mean firing rate via external stimuli or modulating drives tends to reduce neural variability.

**Author Summary:** A topic of considerable current interest concerns the neural mechanisms underlying the suppression of cortical variability following the onset of a stimulus. Since trial-by-trial variability and noise correlations are known to affect the information capacity of neurons, such suppression could improve the accuracy of population codes. One of the main candidate mechanisms is the suppression of noise-induced transitions between multiple attractors, as exemplified by ring attractor networks. The latter have been used to model experimentally measured stochastic tuning curves of directionally selective middle temporal (MT) neurons. In this paper we show how the stimulus-dependent tuning of neural variability in ring attractor networks can be analyzed in terms of the stochastic wandering of spontaneously formed tuning curves or bumps in a continuum neural field model. The advantage of neural fields is that one can derive explicit mathematical expressions for the second-order statistics of neural activity, and explore how this depends on important model parameters, such as the level of noise, the strength of recurrent connections, and the input contrast.

## Introduction

A growing number of experimental studies have investigated neural variability across a variety of cortical areas, brain states and stimulus conditions [6, 21, 23, 35, 45, 51, 52, 58, 59, 63, 72]. Two common ways to measure neural variability are the Fano factor, which is the ratio of the variance to the mean of the neural spike counts over trials, and the trial-to-trial covariance of activity between two simultaneously recorded neuron. It is typically found that the presentation of a stimulus reduces neural variability [21, 51], as does attention and perceptual learning [23, 58]. Another significant feature of the stimulus-dependent suppression of neural variability is that it can be tuned to different stimulus features. In particular, Ponce-Alvarez et al [63] examined the *in vivo* statistical responses of direction selective area-middle temporal (MT) neurons to moving gratings and plaid patterns. They determined the baseline levels and the evoked directional and contrast tuning of the variance of individual neurons and the noise correlations between pairs of neurons with similar direction preferences. The authors also computationally explored the effect of an applied stimulus on variability and correlations in a stochastic ring network model of direction selectivity. They found experimentally that both the trial-by-trial variability and the noise correlations among MT neurons were suppressed by an external stimulus and exhibited bimodal directional tuning. Moreover, these results could be reproduced in a stochastic ring model, provided that the latter operated close to or beyond the bifurcation point for the existence of spontaneous bump solutions.

From a theoretical perspective, a number of different dynamical mechanisms have been proposed to explain aspects of stimulus-dependent variability: (i) stimulus-induced suppression of noise-induced transitions between multiple attractors as exemplified by the stochastic ring model [25, 26, 55, 60, 63]; (ii) stimulus-induced suppression of an otherwise chaotic state [1, 65]; (iii) fluctuations about a single, stimulus-driven attractor in a stochastic stabilized supralinear network [40]. The pros and cons of the different mechanisms have been explored in some detail within the context of orientation selective cells in primary visual cortex (V1) [40]. We suspect that each of the three mechanisms may occur, depending on the particular operating conditions and the specific cortical area. However, we do not attempt to differentiate between these distinct mechanisms in this paper. Instead, we focus on the attractor-based mechanism considered by Ponce-Alvarez et al [63], in order to understand the stimulus-dependent variability of population tuning curves. Our main goal is to show how the tuning of neural variability can be analyzed in terms of the stochastic wandering of spontaneously formed tuning curves or bumps in a continuum neural field model. (For complementary work on the analysis of wandering bumps within the context of working memory see Refs. [46, 47, 49].) The advantage of using neural field theory is that one can derive explicit mathematical expressions for the second-order statistics of neural activity, and explore how this depends on important model parameters, such as the level of noise, the strength of recurrent connections, and the input contrast. In particular, our mathematical analysis provides a simple explanation for the bimodal tuning of the variance observed by Ponce-Alvarez et al [63].

In addition to accounting for the qualitative statistical behavior of a single ring network, we also explore the effects of inter-network coupling on stimulus-dependent variability in a pair of ring networks. The latter could represent populations of cells in two different layers of a cortical hypercolumn linked via vertical synaptic connections, or two different cortical hypercolumns linked by horizontal patchy connections within the same layer. We will refer to these two distinct architectures as model A and model B, respectively. (See also Figs. 14 and 2.) Roughy speaking, cortical layers can be grouped into input layer 4, superficial layers 2/3 and deep layers 5/6 [16, 28, 42, 62]. They can be distinguished by the source of afferents into the layer and the targets of efferents leaving the layer, the nature and extent of intralaminar connections, the identity of interneurons within and between layers, and the degree of stimulus specificity of pyramidal cells. In previous work, we explored the role of cortical layers in the propagation of waves of orientation selectivity across V1 [13]. In this paper, we use model A to show how vertical excitatory connections between two stochastic ring networks can reduce neural variability, consistent with a previous analysis of spatial working memory [47]. We also show that the degree of noise suppression can differ between layers, as previously found in an experimental study of orientation selective cells in V1 [74]. Long-range horizontal connections within superficial layers of cortex are mediated by the axons of excitatory pyramidal neurons. However, they innervate both pyramidal neurons and feedforward interneurons so that they can have a net excitatory or inhibitory effect, depending on stimulus conditions [3, 4, 71]. More specifically, they tend to be excitatory at low contrasts and inhibitory at high contrasts. An experimental “center-surround” study of stimulus-dependent variability in V1 indicates that correlations in spontaneous activity at the center can be suppressed by stimulating outside the classical receptive field of the recorded neurons [75], that is, by evoking activity in the surround. In this paper, we show that the effect of a surround stimulus depends on at least two factors: (i) whether or not the horizontal connections effectively excite or inhibit the neurons in the center, and (ii) the relative directional bias of the surround stimulus. In particular, we find that at low contrasts (excitatory regime), noise is suppressed in the center when the center and surround stimuli have the same directional bias, whereas it is facilitated when the center and surround stimuli have opposite directional biases. The converse holds at high contrasts (inhibitory regime). These results are consistent with the general observation that increasing the mean firing rate via external stimuli or modulating drives tends to reduce neural variability.

In the remainder of the **Introduction** we introduce our stochastic neural field model of coupled ring networks and describe in more detail the structure of models A and B. We then briefly summarize previous work on the theory of wandering bumps. In **Materials and Methods** we use perturbation theory to show how the neural field equations can be reduced to a pair of stochastic phase equations describing the stochastic wandering of bump solutions. These equations are analyzed in the **Results**, using a modified version of the bivariate von Mises distribution, which is well-known in the theory of circular statistics. This then allows us to determine the second-order statistics of a single ring network, providing a mathematical underpinning for the experimental and computational studies of Ponce-Alvarez et al [63], and to explore the effects of inter-network coupling on neural variability in models A and B.

## Coupled ring model

Consider a pair of mutually coupled ring networks labeled *j* = 1, 2. Let *u*_*j*_ (*θ*, *t*) denote the activity at time *t* of a local population of cells with stimulus preference *θ* ∈ [−*π*, *π*) in network *j*. Here *θ* could represent the direction preference of neurons in area-middle temporal cortex (MT) [63], the orientation preference of V1 neurons, after rescaling *θ* → *θ*/2 [8, 9], or a coordinate in spatial working memory [17, 47, 54]. For concreteness, we will refer to *θ* as a direction preference. The variables *u*_*j*_ evolve according to the neural field equations [10, 46, 47, 83]

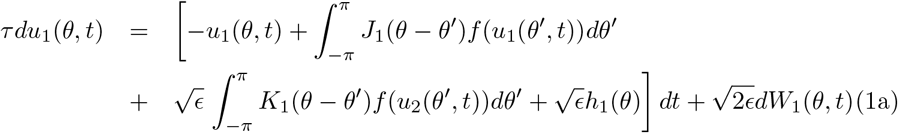

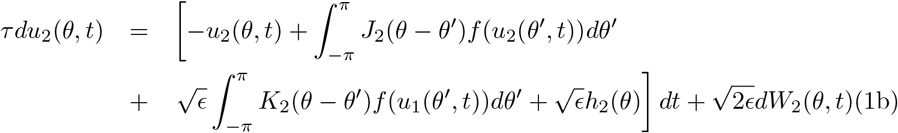

where 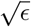 is a constant scale factor (see below), *J*_*j*_ (*θ* − *θ*′) is the distribution of intra-network connections between cells with stimulus preferences *θ*′ and *θ* in network *j*, *K*_*j*_ (*θ* − *θ*′) is the corresponding distribution of inter-network connections to network *j*, and *h*_*j*_ (*θ*) is an external stimulus. The firing rate function is assumed to be a sigmoid

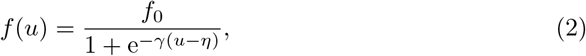

with maximal firing rate *f*_0_, gain *γ* and threshold *η*. The final term on the right-hand side of each equation represents external additive noise, with *W*_*j*_ (*θ*, *t*) a *θ*-dependent Wiener process. In particular,

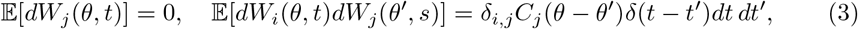

where *δ*(*t*) is the Dirac delta function. The noise is thus colored in *θ* (which is necessary for the solution to be spatially continuous) and white in time. (One could also take the noise to be colored in time by introducing an additional Ornstein-Uhlenbeck process. For simplicity, we assume that the noise processes in the two networks are uncorrelated, which would be the case if the noise were predominantly intrinsic. Correlations would arise if some of the noise arose from shared fluctuating inputs. For a discussion of the effects of correlated noise in coupled ring networks see [47].) The external stimuli are taken to be weakly biased inputs of the form

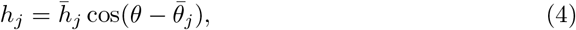

where 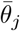 is the location of the peak of the input (stimulus bias) and 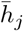 is the contrast. Finally, the time-scale is fixed by setting the time constant *τ* = 10 msec. The maximal firing rate *f*_0_ varies between 10-100 spikes/sec.

The weight distributions are 2*π*-periodic and even functions of *θ* and thus have cosine series expansions. Following [46], we take the intra-network recurrent connections to be

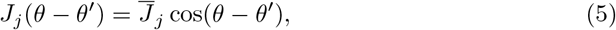

which means that cells with similar stimulus preferences excite each other, whereas those with sufficiently different stimulus preferences inhibit each other. It remains to specify the nature of the inter-network connections. As we have already mentioned, we consider two different network configurations: (A) a vertically connected two layer or laminar model (Fig. 14) and (B) a horizontally connected single layer model (Fig. 2). In model A, the inter-network weight distribution is taken to have the form

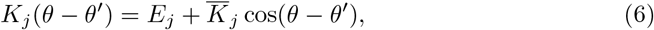

which represents vertical coupling between the layers. We also assume that both layers receive the same stimulus bias, that is, 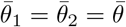 in equation (4). In model B, the inter-network weight distribution represents patchy horizontal connections, which tend to link cells with similar stimulus preferences. This is implemented by taking

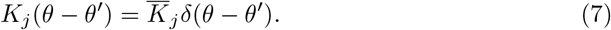

**Figure 1.**
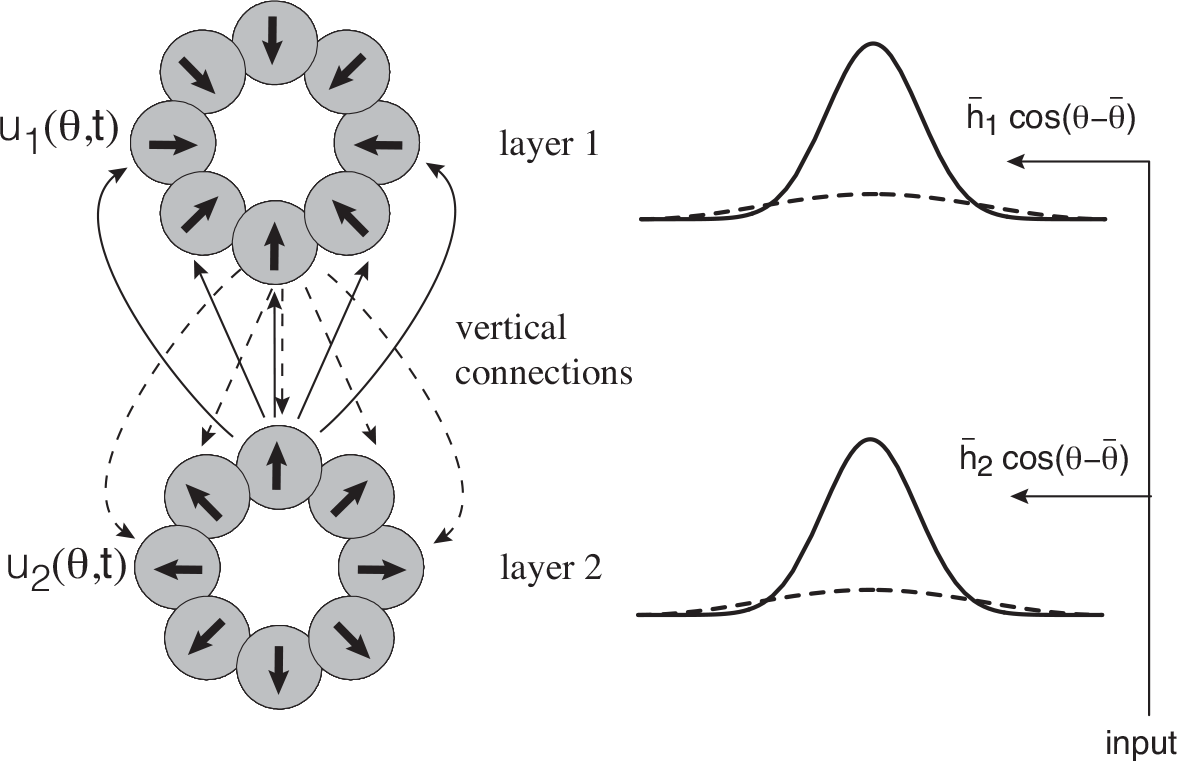
Coupled ring model A: the two ring networks are located in two vertically separated cortical layers and interact via interlaminar connections

Now the two networks can be driven by stimuli with different biases so that 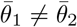.

Note that in order to develop the analytical methods of this paper, we scale the internetwork coupling, the noise terms and the external stimuli in equations (1) by the constant factor 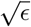. Taking 0 < *ϵ* ≪ 1 (weak noise, weak inputs and weak inter-network coupling) will allow us to use perturbation methods to derive explicit parameter-dependent expressions for neural variability. We do not claim that cortical networks necessarily operate in these regimes, but use the weakness assumption to obtain analytical insights and make predictions about the qualitative behavior of neural variability. In the case of weak inter-network connections, the validity of the assumption is likely to depend on the source of these connections. For example, in model B, they arise from patchy horizontal connections within superficial or deep layers of cortex, which are known to play a modulatory role [4]. On the other hand, vertical connections between layers are likely to be stronger than assumed in our modeling analysis, at least in the feedforward direction.

**Figure 2.**
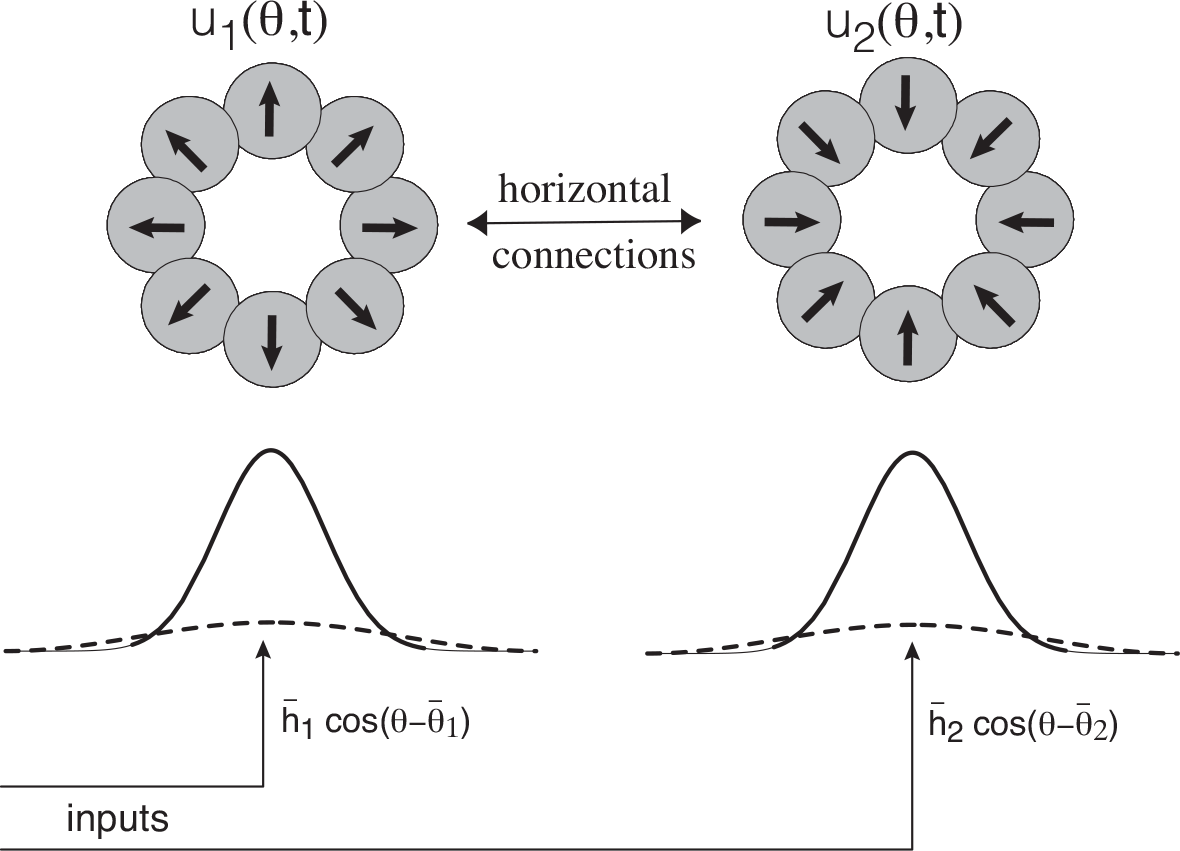
Coupled ring model B: the two ring networks are located in the same cortical layer and interact via intralaminar horizontal connections.

The weak stimulus assumption depends on a particular view of how cortical neurons are tuned to stimuli. Consider the most studied example, which involves orientation tuning of cells in V1. The degree to which recurrent processes contribute to the receptive field properties of V1 neurons has been quite controversial over the years [33, 64, 77, 82]. The classical model of Hubel and Wiesel [43] proposed that the orientation preference and selectivity of a cortical neuron in input layer 4 arises primarily from the geometric alignment of the receptive fields of thalamic neurons in the lateral geniculate nucleus (LGN) projecting to it. (Orientation selectivity is then carried to other cortical layers through vertical projections). This has been confirmed by a number of experiments [32, 34, 66, 69, 79]. However, there is also significant experimental evidence suggesting the importance of recurrent cortical interactions in orientation tuning [29, 50, 61, 68, 70, 73, 78]. One issue that is not disputed is that some form of inhibition is required to explain features such as contrast-invariant tuning curves and cross-orientation suppression [64]. The uncertainty in the degree to which intracortical connections contribute to orientation tuning of V1 neurons is also reflected in the variety of models. In ring attractor models [7, 9, 76], the width of orientation tuning of V1 cells is determined by the lateral extent of intracortical connections. Recurrent excitatory connections amplify weakly biased feedforward inputs in a way that is sculpted by lateral inhibitory connections. Hence, the tuning width and other aspects of cortical responses are primarily determined by intracortical rather than thalamocortical interconnections. On the other hand, in push-pull models, cross-orientation inhibition arises from feedforward inhibition from interneurons [79, 80]. Finally, in normalization models, a large pool of orientation-selective cortical interneurons generates shunting inhibition proportional in strength to the stimulus contrast at all orientations [18]. In the end, it is quite possible that are multiple circuit mechanisms for generating tuned cortical responses to stimuli, which could depend on the particular stimulus feature, location within a feature preference map, and cortical layer [64]. In our analytical study we adopt the ring attractor network model, for which inputs are weak.

## Wandering bumps and neural variability

There have been numerous previous studies establishing that a single homogeneous neural field on the real line or a ring can support a stationary bump or population tuning curve in the absence of external stimuli [2, 7–9, 46, 53, 76, 81, 84]. The network is said to be in a marginally stable regime, since the location of the peak of the bump is arbitrary. This reflects the fact that the homogeneous neural field is symmetric with respect to uniform translations. The presence of a weakly biased external stimulus can then lock the bump to the stimulus. The output activity is said to amplify the input bias and provides a network-based encoding of the stimulus, which can be processed by upstream networks. Since the bump persists if the stimulus is removed, marginally stable neural fields have been proposed as one mechanism for implementing a form of spatial working memory [17, 20, 47, 54, 85]. One of the consequences of operating in a marginally stable regime is that the bump is not robust to the effects of external noise, which can illicit a stochastic wandering of the bump [19, 20, 46, 47, 53].

One way to investigate the stochastic wandering of bumps in a neural field model is to use perturbation theory. The latter was originally applied to the analysis of traveling waves in one-dimensional neural fields [10, 83], and was subsequently extended to the case of wandering bumps in single-layer and multi-layer neural fields [14, 46, 47]. The basic idea is to to treat longitudinal and transverse fluctuations of a bump (or traveling wave) separately in the presence of noise, in order to take proper account of marginal stability. This is implemented by decomposing the stochastic neural field into a deterministic bump profile, whose spatial location or phase has a slowly diffusing component, and a small error term. (There is always a non-zero probability of large deviations from the bump solution, but these are assumed to be negligible up to some exponentially long time.) Perturbation theory can then be used to derive an explicit stochastic differential equation (SDE) for the diffusive-like wandering of the bump in the weak noise regime. (A more rigorous mathematical treatment that provides bounds on the size of transverse fluctuations has also been developed [15, 44].) In this paper, we apply the theory of wandering bumps in stochastic neural fields in order to characterize various features of stimulus-dependent variability in cortical neurons.

## Results

We present various analytical and numerical results concerning stimulus-dependent neural variability, under the assumption that the neural field equations (1) support stable stationary bump solutions *u*_*j*_ (*θ*, *t*) = *U*_*j*_ (*θ*) = *A*_*j*_ cos(*θ*), *j* = 1, 2, in the absence of noise, external stimuli, and inter-network coupling (*ϵ* = 0). The amplitudes *A*_*j*_ are determined self-consistently from the equations (see **Material and Methods**)

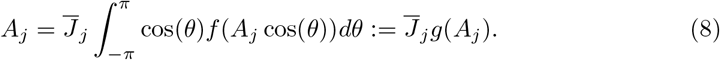

In order to investigate the effects of noise in the presence of weak external stimuli and inter-network coupling, we introduce the amplitude phase decomposition [10, 47]

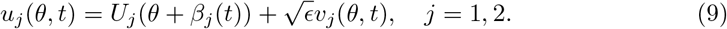

Such a decomposition reflects the fact that the uncoupled homogeneous neural field equations are marginally stable with respect to uniform translations around the ring. Substituting equations (9) into the full stochastic neural field equations (1) and using perturbation theory along the lines of [10, 14, 46, 47], one can derive the following SDEs for the phases *β*_*j*_ (*t*), see **Materials and Methods**:

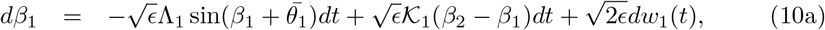

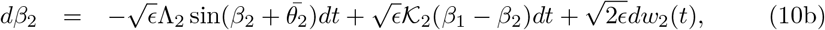

where 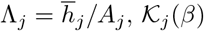 are 2*π*-periodic functions that depend on the form of the inter-network connections, and *w*_*j*_ (*t*) are independent Wiener processes:

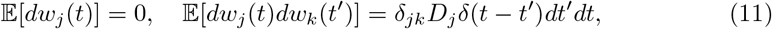

The functions 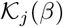 and the diffusion coefficients *D*_1_, *D*_2_ are calculated in **Materials and Methods**, see equations (72), (73) and (76).

Note that stochastic phase equations similar to (10) were previously derived in [46, 47], except that sin *β* and 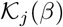 were linearized (eg. sin *β ≈ β* etc.), resulting in a system of coupled Ornstein-Uhlenbeck (OU) processes. Properties of one-dimensional OU processes were then used to explore how the variance in the position of bump solutions depended on inter-network connections and statistical noise correlations. However, it should be noted that the variables *β*_*j*_ (*t*) are phases on a circle (rather than positions on the real line), so that the right-hand side of equations (10) should involve 2*π*-period functions. Therefore, the linear approximation only remains accurate on sufficiently short times scales for which the probability of either of the phases winding around the circle is negligible. Here we show how to analyze the full nonlinear phase equations, and use this to explore the effects of external stimuli and inter-network connections on neural variability.

### Wandering bumps in a single stochastic ring network

Let us begin by considering stimulus-dependent neural variability in a single ring network evolving according to the stochastic neural field equation

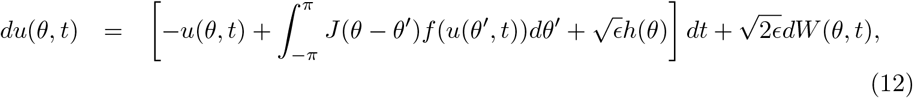

where

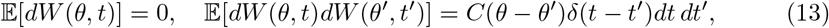

with

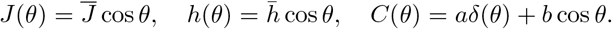

A clear demonstration of the suppressive effects of an external stimulus can be seen from direct numerical simulations of equation (12), see Figs. 3 and 4. In the absence of an external stimulus, the center-of-mass (phase) of the bump diffuses on the ring, whereas it exhibits localized fluctuations when a weakly-biased stimulus is present.

**Figure 3.**
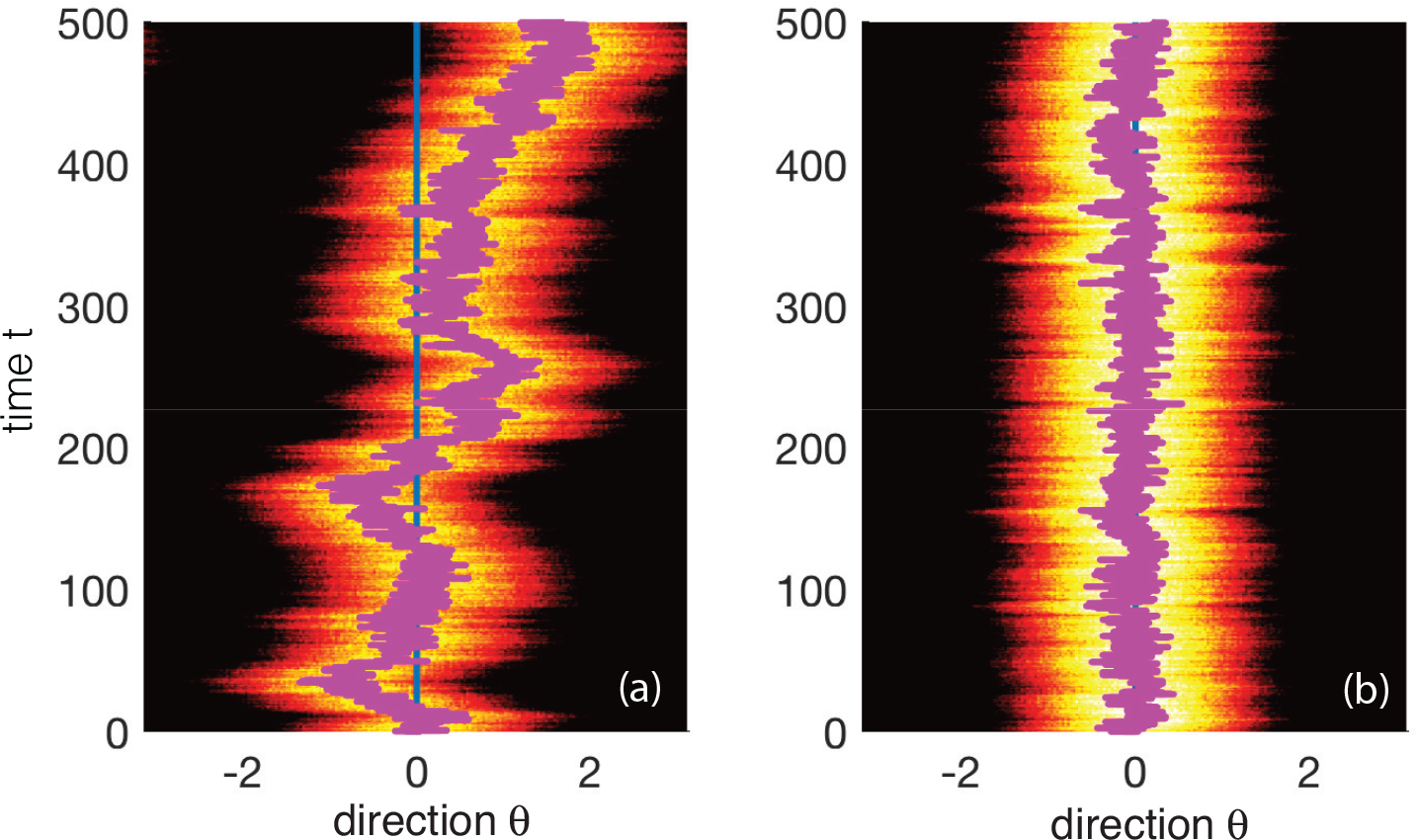
Stimulus-dependent wandering of a bump in a single stochastic ring network. Overlaid lines represent the trajectory of the center-of-mass or phase of the bump, *β*(*t*). (a) In the absence of an external stimulus 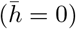, the center-of-mass of the bump executes diffusive-like motion on the ring. (b) The presence of a weakly biased external stimulus 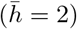 significantly suppresses fluctuations, localizing the bump to the stimulus direction 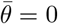. Parameters are threshold *η* = 0.5, gain *γ* = 4, synaptic weight 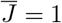, correlation parameters *a* = 3, *b* = 0.5 and *ϵ* = 0.05.

**Figure 4.**
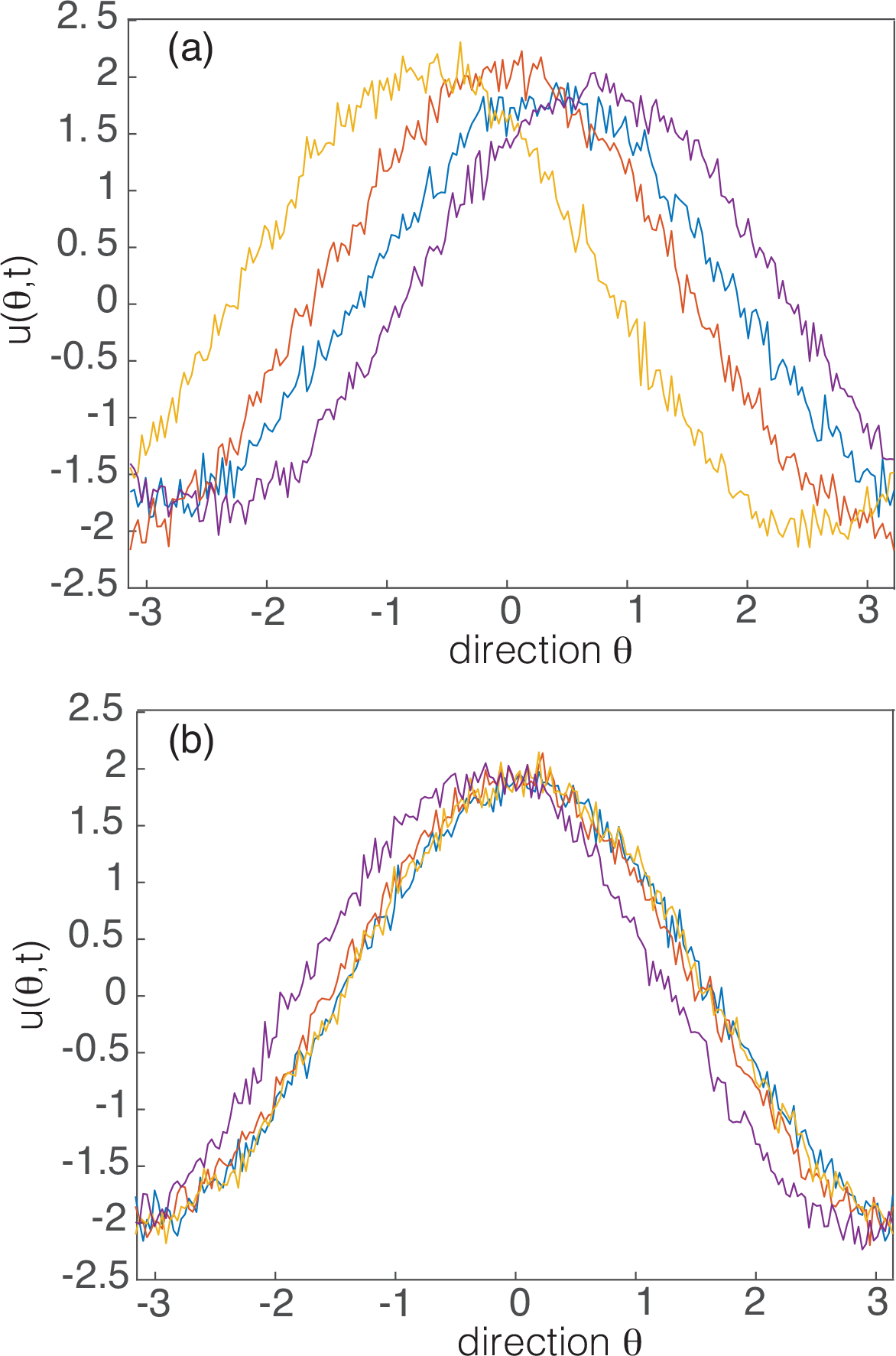
Snapshots of bump profiles at different times. (a) For no external stimulus the bumps are distributed at different positions around the ring and vary in amplitude. (b) In the presence of a stimulus the bumps are localized around zero and have similar amplitudes.

Clearly, the main source of neural variation is due to the wandering of the bump, which motivates the amplitude phase decomposition given by equation (9).

Applying the perturbation analysis of **Materials and Methods** yields a one-network version of the phase equations (10), which takes the form

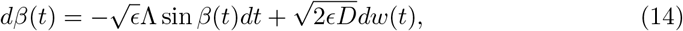

with 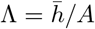 and 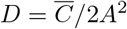, where *A* is the amplitude of the bump for *ϵ* = 0. Equation (14) is known as a von Mises process, which can be regarded as a circular analog of the Ornstein-Uhlenbeck process on a line, and generates distributions that frequently arise in circular or directional statistics [56]. The von Mises process has been used to model the trajectories of swimming organisms [22, 41], oscillators in physics [38], bioinformatics [57], and the data fitting of neural population tuning curves [5]. (Nonlinear stochastic phase equations analogous to (14) also arise in models of ring attractor networks with synaptic heterogeneities, which have applications to spatial working memory [48, 49, 67].)

Introduce the probability density

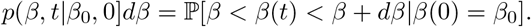

This satisfies the forward Fokker-Planck equation (dropping the explicit dependence on initial conditions)

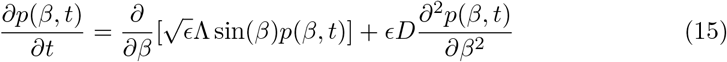

for *β* ∈ [-*π, π*] with periodic boundary conditions *p*(−*π, t*) = *p*(*π, t*). It is straightforward to show that the steady-state solution of equation (15) is the von Mises distribution

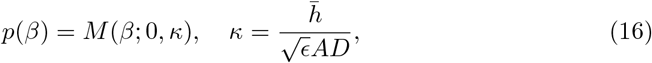

with

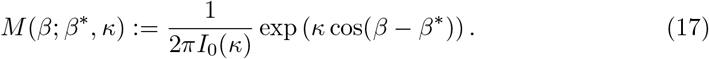

Here *I*_0_(*κ*) is the modified Bessel function of the first kind and zeroth order (*n* = 0), where

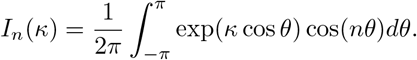

Sample plots of the von Mises distribution are shown in Fig. 5. One finds that *M*(*β*; *β*\*, *κ*) → 1/2*π* as *κ* → 0; since 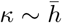 this recovers the uniform distribution of pure Brownian motion on the circle. On the other hand, the von Mises distribution becomes sharply peaked as *κ* → ∞. More specifically, for large positive *κ*,

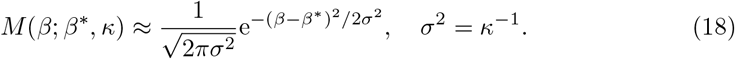

**Figure 5.**
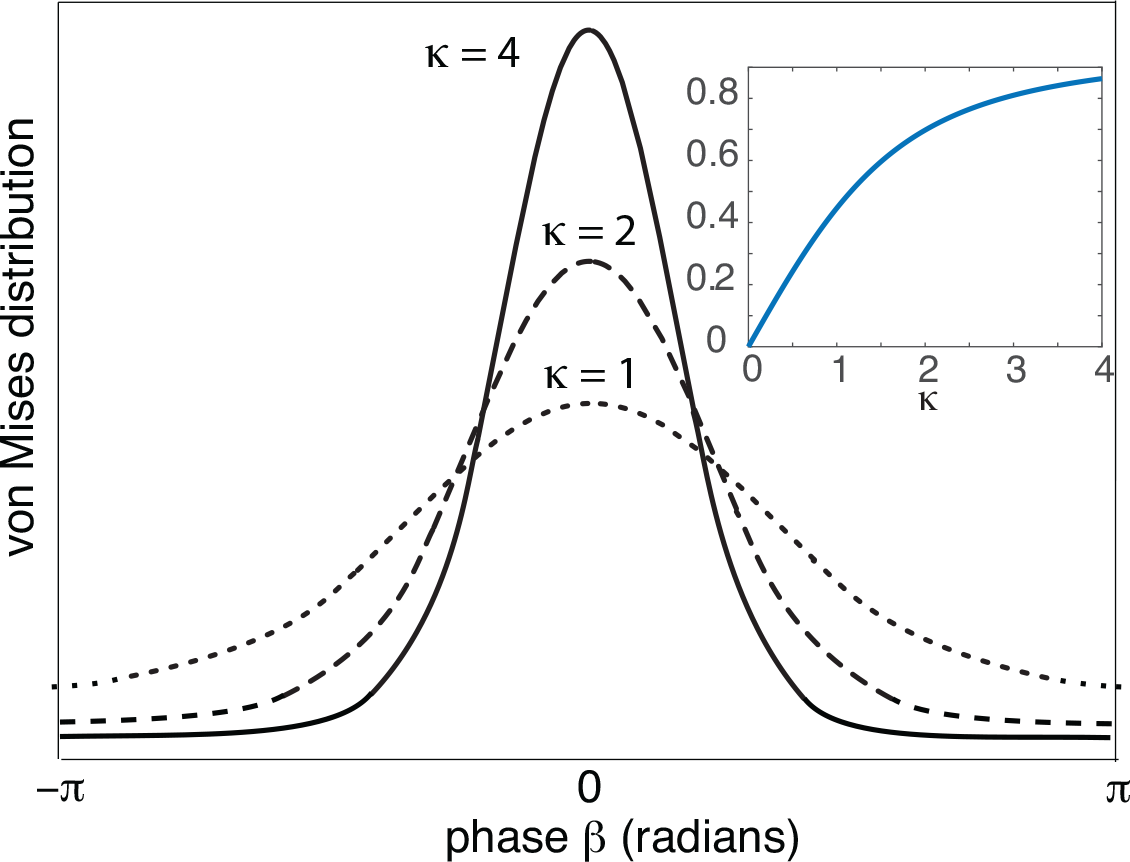
Sample plots of the von Mises distribution *M*(*β,* 0, *κ*) centered at zero for various values of *κ*. Inset: Plot of first circular moment *I*_1_(*κ*)/*I*_0_(*κ*).

We thus have an explicit example of the noise suppression of fluctuations by an external stimulus, since 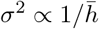.

Moments of the von Mises distribution are usually calculated in terms of the circular moments of the complex exponential *x* = e^*iβ*^ = cos *β* + *i* sin *β*. The *n*th circular moment is defined according to

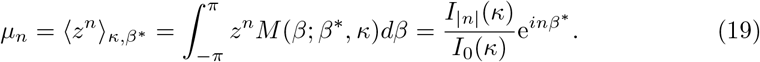

In particular,

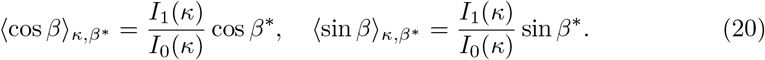

We can use these moments to explore stimulus-dependent variability in terms of the stochastic wandering of the bump or tuning curve. That is, consider the leading order approximation *u*(*θ, t*) ≈ *A* cos(*θ* + *β*(*t*)), with *β*(*t*) evolving according to the von Mises SDE (14). Trial-to-trial variability can be captured by averaging the solution with respect to the stationary von Mises density (16). First,

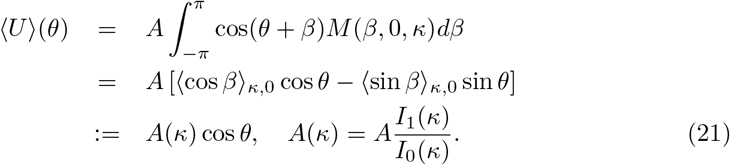

from equations (20). Hence, the mean amplitude *A*(*κ*) is given by the first circular moment of the von Mises distribution, see inset of Fig. 5. When *κ* = 0 (zero external stimulus), the amplitude vanishes due to the fact that the random position of the bump is uniformly distributed around the ring. As the stimulus contrast 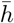 increases the wandering of the bump is more restricted and *A*(*κ*) monotonically increases.

Second,

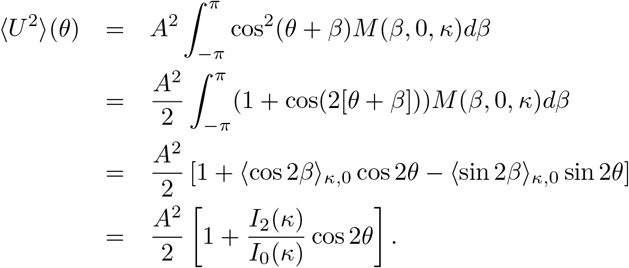

It follows that the variance is

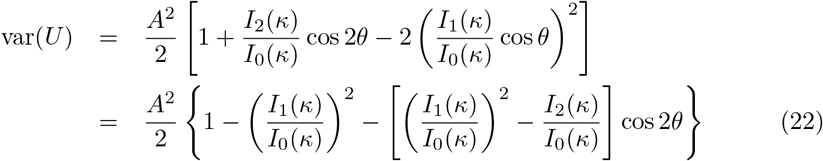

**Figure 6.**
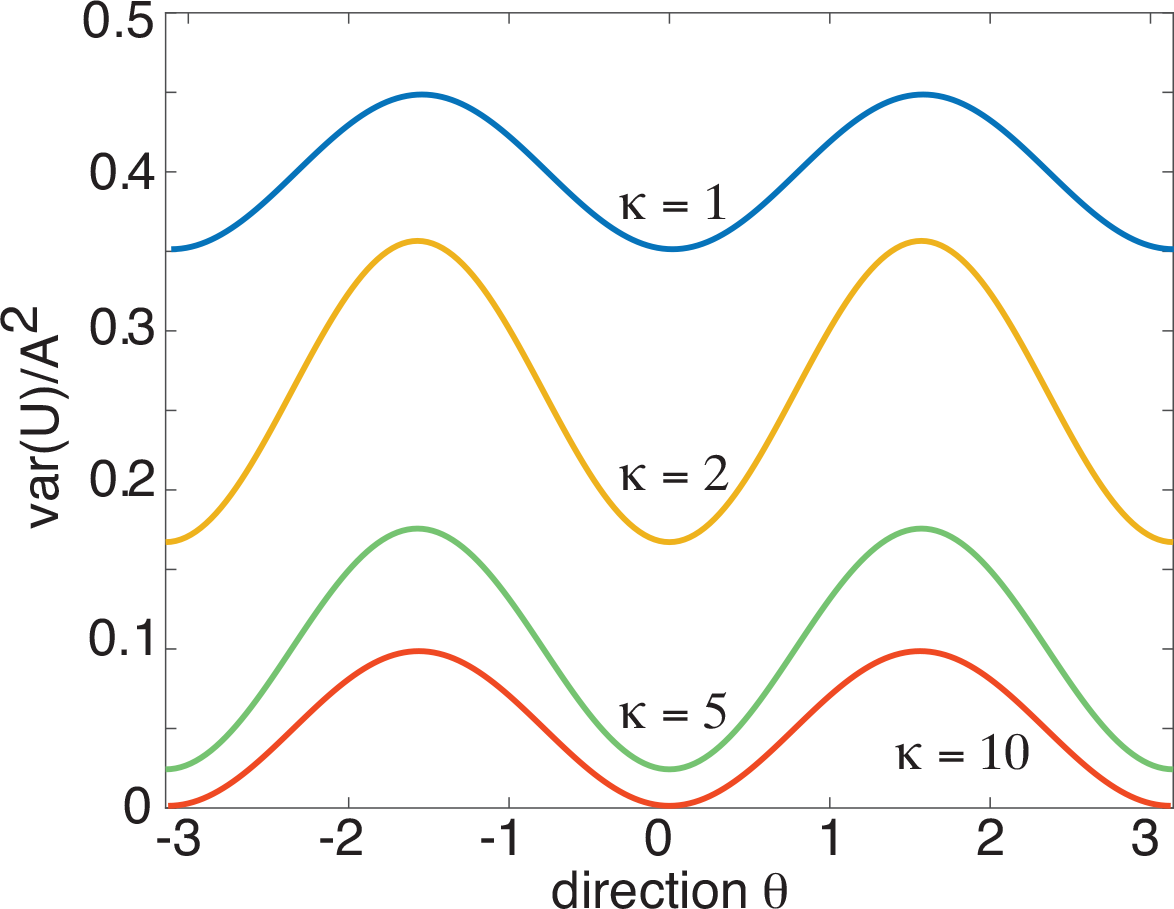
Plot of normalized variance var(*U*)*/A*^2^ as a function of *θ* for a single ring network and various *κ*. In the spontaneous case (*κ* = 0) the variance is uniformly distributed around the ring (ignoring transients). The presence of a stimulus (*κ* > 0) suppresses the overall level of noise and the variance exhibits a bimodal tuning curve.

In Fig. 6, we show example plots of the normalized variance var(*U*)*/A*^2^ as a function of the parameter *κ*, which is a proxy for the input amplitude 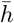, since 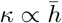. It can be seen that our theoretical analysis reproduces the various trends observed in [63]: (i) a global suppression of neural variability that increases with the stimulus contrast; (ii) a directional tuning of the variability that is bimodal; (iii) a peak in the suppression of cells at the preferred directional selectivity. One difference between our theoretical results and those of [63] is that, in the latter case, the directional tuning of the variance is not purely sinusoidal. Part of this can be accounted for by noting that we consider the variance of the activity variable *u* rather than the firing rate *f*(*u*). Moreover, for analytical convenience, we take the synaptic weight functions etc. to be first-order harmonics. In Fig. 7 we show numerical plots of the variance in the firing rate, which exhibits the type of bimodal behavior found in [63] when the ring network operates in the marginal regime.

**Figure 7.**
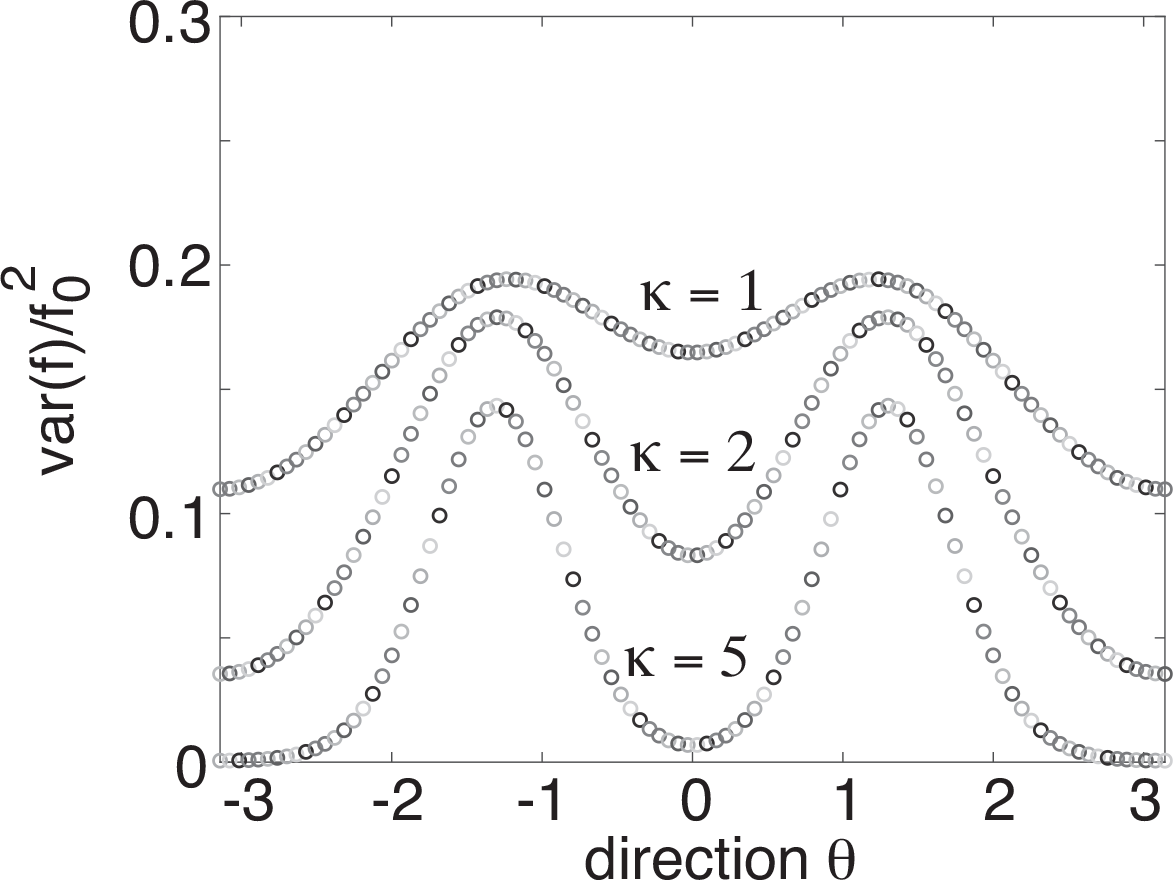
Plot of variance in firing rates var(*f*(*A* cos(*θ*)) (in units of 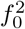) as a function of *θ* for a single ring network and various *κ*. *f* is given by the sigmoid function (2) with *γ* = 4 and *η* = 0.5. The corresponding amplitude *A* ≈ 1.85.

### Effects of inter-network coupling (model A)

We now turn to a pair of coupled ring networks that represent vertically connected layers as shown in Fig. 14 (model A), with inter-network weight distribution (6). For analytical tractability, we impose the symmetry conditions *A*_1_ = *A*_2_ = *A* and 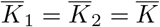. However, we allow the contrasts of the external stimuli to differ, 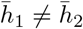. Also, without loss of generality, we set 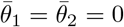. Equations (10) then reduce to the form, see **Materials and Methods**

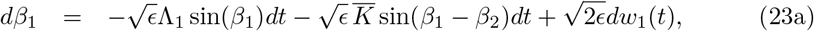

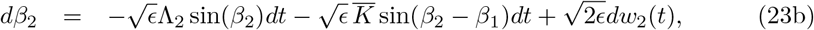

with

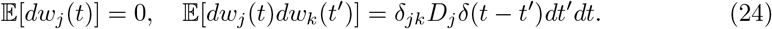

Given our various simplifications, we can rewrite equations (23) in the more compact form

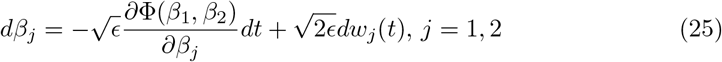

where Φ is the potential function

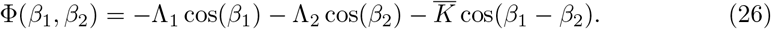

Introduce the joint probability density

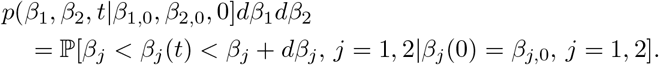

This satisfies the two-dimensional forward Fokker-Planck equation (dropping the explicit dependence on initial conditions)

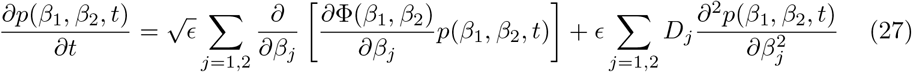

for *β*_*j*_ ∈ [−*π, π*] and periodic boundary conditions *p*(−*π*, *β*_2_, *t*) = *p*(*π*, *β*_2_, *t*), *p*(*β*_1_, −*π*, *t*) = *p*(*β*_1_, *π*, *t*).

The existence of a potential function means that we can solve the time-independent FP equation. Setting time derivatives to zero, we have

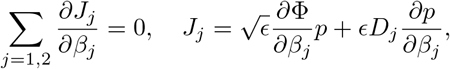

where *J*_*j*_ is a probability current. In the stationary state the probability currents are constant, but generally non-zero. However, in the special case *D*_1_ = *D*_2_ = *D*, then there exists a steady-state solution in which the currents vanish. This can be seen by rewriting the vanishing current conditions as

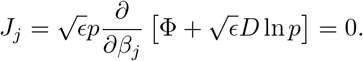

This yields the steady-state probability density, which is a generalization of the von Mises distribution,

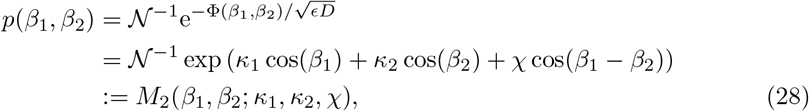

where

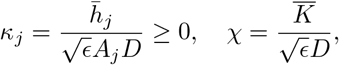

and 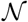 is the normalization factor

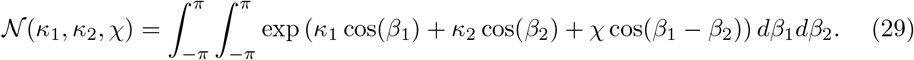

The distribution *M*_2_(*β*_1_, *β*_2_; *κ*_1_, *κ*_2_, *χ*) is an example of a bivariate von Mises distribution known as the cosine model [57]. The normalization factor can be calculated explicitly to give

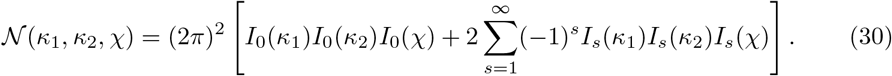

The corresponding marginal distribution for *β*_1_ is

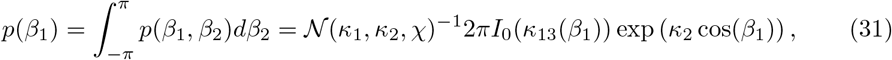

where

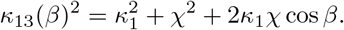

An analogous result holds for the marginal density *p*(*β*_2_).

We now summarize a few important properties of the cosine bivariate von Mises distribution [57]:

1. The density *M*_2_(*β*_1_, *β*_2_; *κ*_1_, *κ*_2_, *χ*) is unimodal if

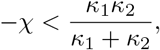

and is bimodal if

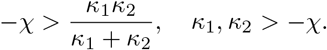
2. When *κ*_1_ and *κ*_2_ are large, the random variables (*β*_1_, *β*_2_) are approximately bivariate normal distributed, that is, (*β*_1_, *β*_2_) ~ *N*_2_(0, Σ) with

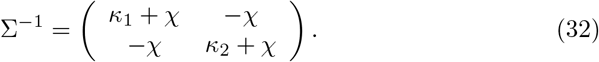

We will assume that the vertical connections are maximal between neurons with the same stimulus preference so that 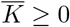 and 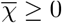. It then follows that *p*(*β*_1_, *β*_2_) is unimodal. Moreover, from equation (32) we have

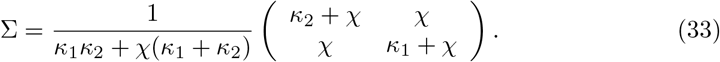

For zero inter-network coupling (*χ* = 0), we obtain the diagonal matrix 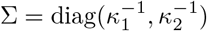 and we recover the variance of the single ring networks, that is, 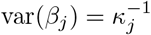; there are no interlaminar correlations. On the other hand, for *χ* > 0 we find two major effects of the interlaminar connections. First, the vertical coupling reduces fluctuations in the phase variables within a layer. This is most easily seen by considering the symmetric case *κ*_1_ = *κ*_2_ = *κ* for which

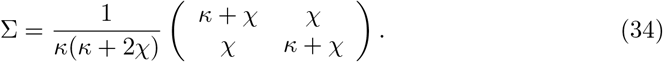

**Figure 8.**
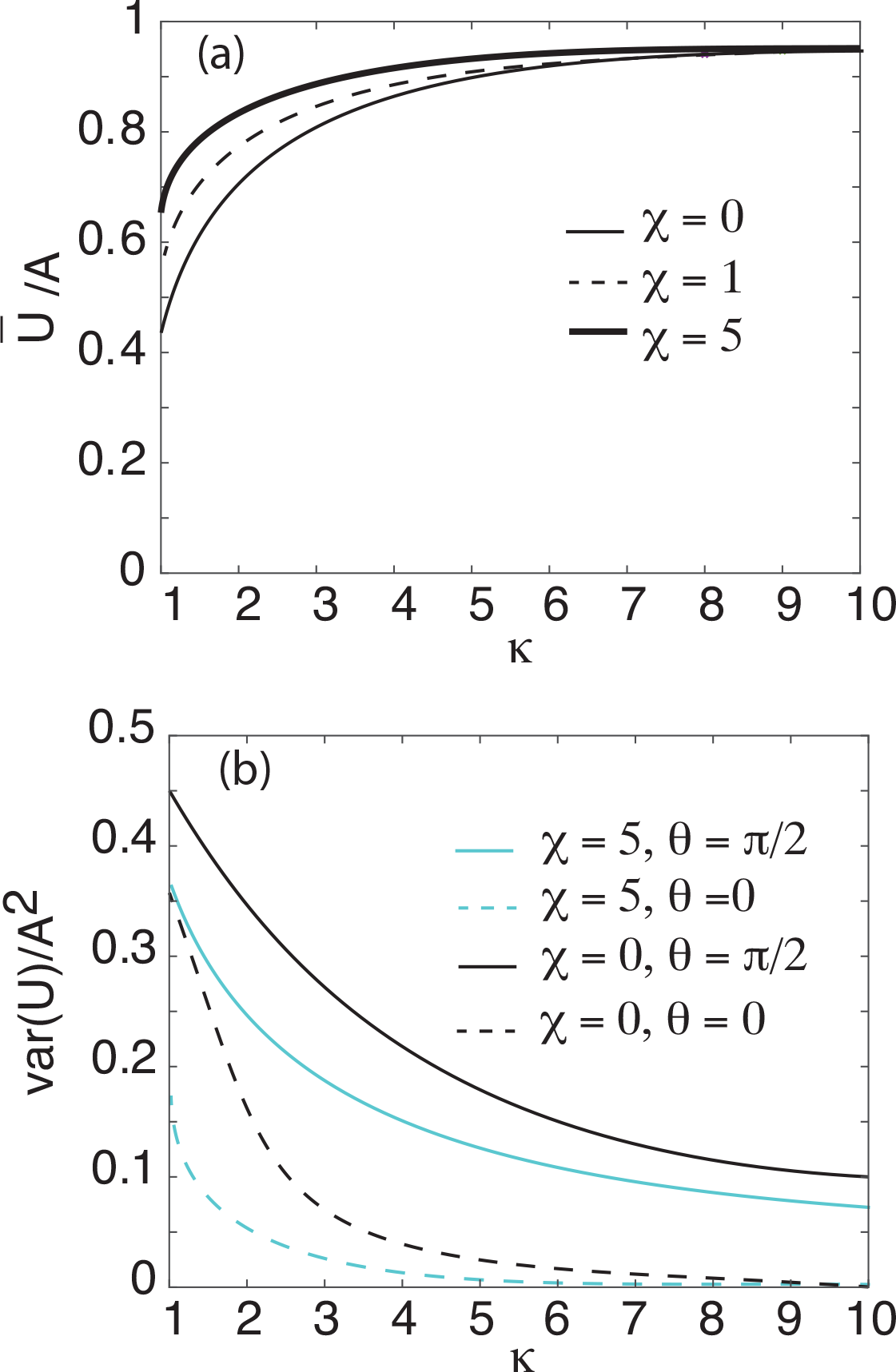
Coupled ring network (model A). (a) Amplitude of normalized mean tuning curve (36) as a function of the input parameter *κ* = *κ*_1_ = *κ*_2_ for various coupling strengths: *χ* = 0, 1, 5. (b) Corresponding maximum (*θ* = π/2) and minimum (*θ* = 0) normalized variances (37) as a function of the input parameter *κ* for coupling strengths *χ* = 0, 5.

**Figure 9.**
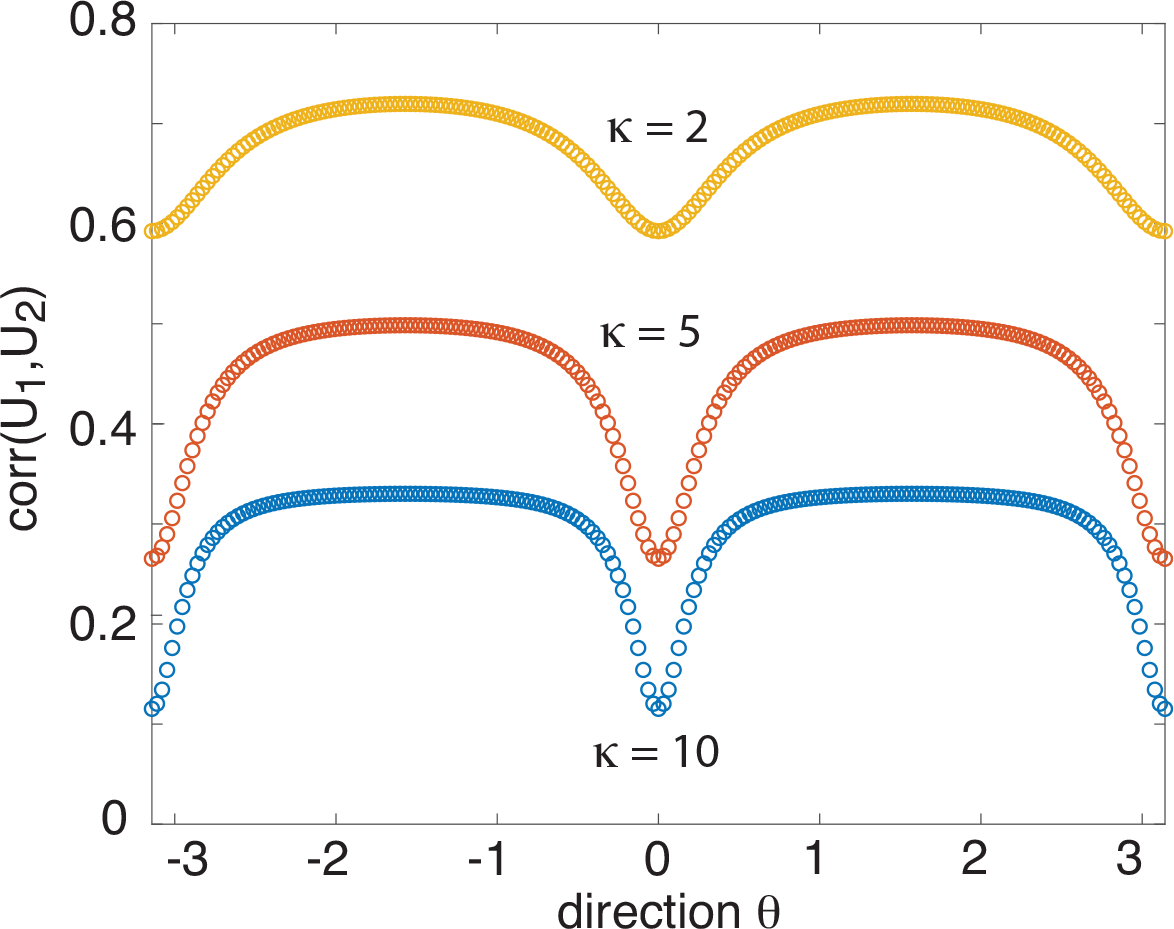
Coupled ring network A. Plot of correlation tuning curve (39) between cells with the same direction preference but located in different layers. Here *κ* = *κ*_1_ = *κ*_2_ and *χ* = 5.

Clearly,

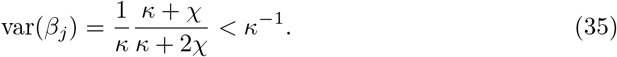

(This result is consistent with a previous study of the effects of inter-network connections on neural variability, which focused on the case of zero stimuli and treated the bump positions as effectively evolving on the real line rather than a circle [47]. In this case, inter-network connections can reduce the variance in bump position, which evolves linearly with respect to the time *t*.) The second consequence of interlaminar connections is that they induce correlations between the phase *β*_1_(*t*) and *β*_2_(*t*).

Having characterized the fluctuations in the phases *β*_1_(*t*) and *β*_2_(*t*), analogous statistical trends will apply to the trial-to-trial variability in the tuning curves. This follows from making the leading-order approximation *u*_*j*_ (*x, t*) ~ *A* cos(*θ* + *β*_*j*_ (*t*)), and then averaging the *β*_*j*_ with respect to the bivariate von Mises density *M*_2_(*β*_1_, *β*_2_; *κ*_1_, *κ*_2_, *χ*). In the large *κ*_*j*_ regime, this could be further simplified by averaging with respect to the bivariate normal distribution under the approximations cos(*β*) ≈ 1 − *β*^2^/2 and sin *β* ~ *β*. Both the mean and variance of the tuning curves are similar to the single ring network, see equations (21) and (22):

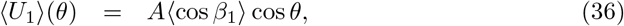

and

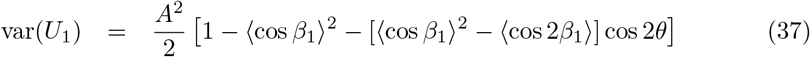

**Figure 10.**
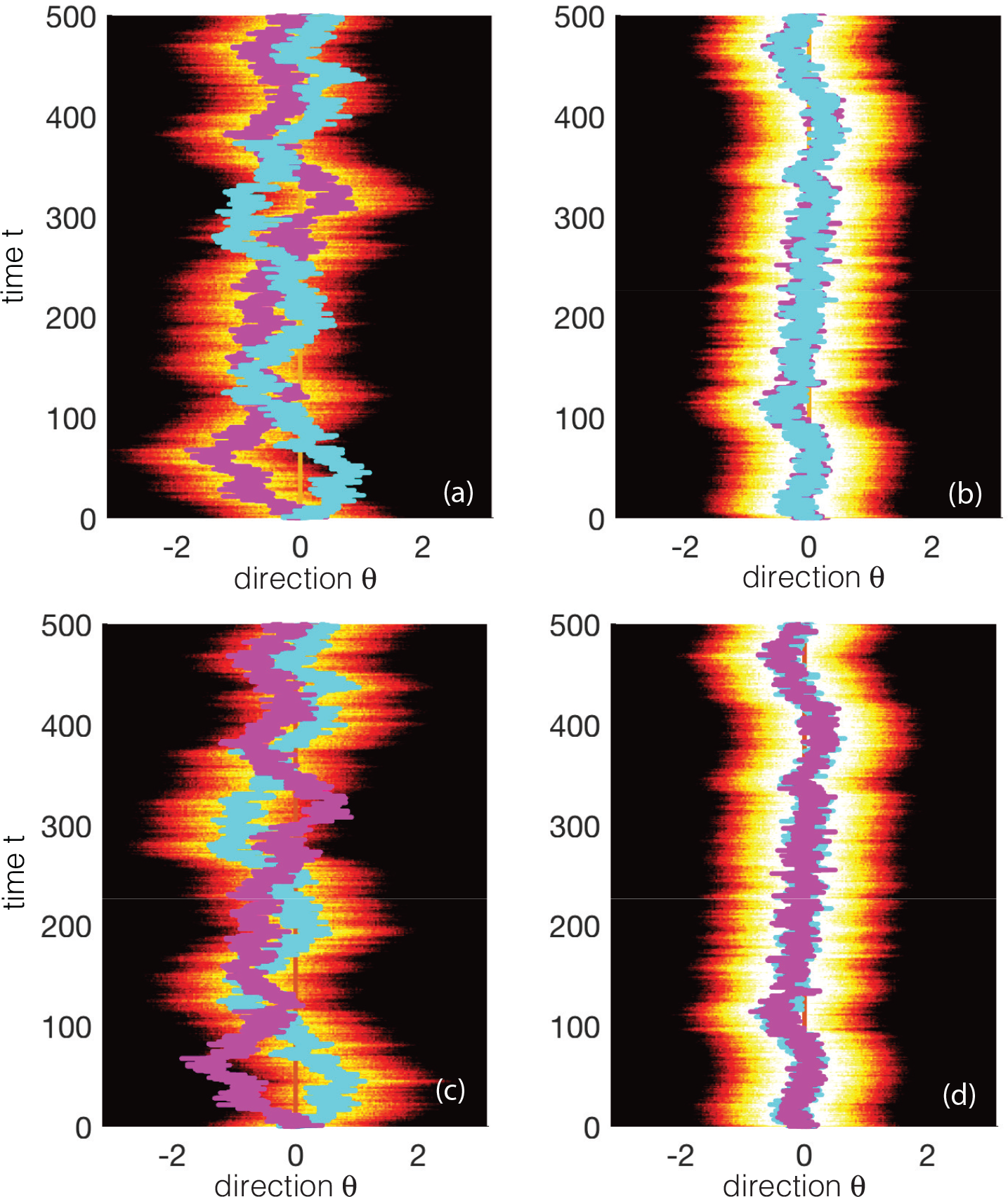
Effects of interlaminar connections on a pair of wandering bumps (model A). Overlaid lines represent the trajectories of the center-of-mass or phase of the bumps, *β*_1_(*t*) and *β*_2_(*t*). (a,b) Plots of wandering bump in network 1 for zero 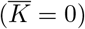 and nonzero 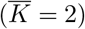 interlaminar connections, respectively. (c,d) Analogous plots for network 2. The two networks are taken to be identical with the same parameters as Fig. 3 except 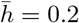.

Their dependence on the coupling strength *χ* and input parameter *κ*_1_ = *κ*_2_ = *κ* is illustrated in Fig. 8. Finally,

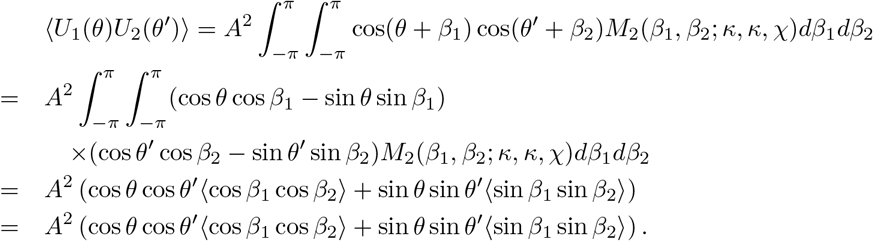

so that inter-network covariance take the form

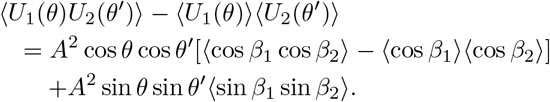

**Figure 11.**
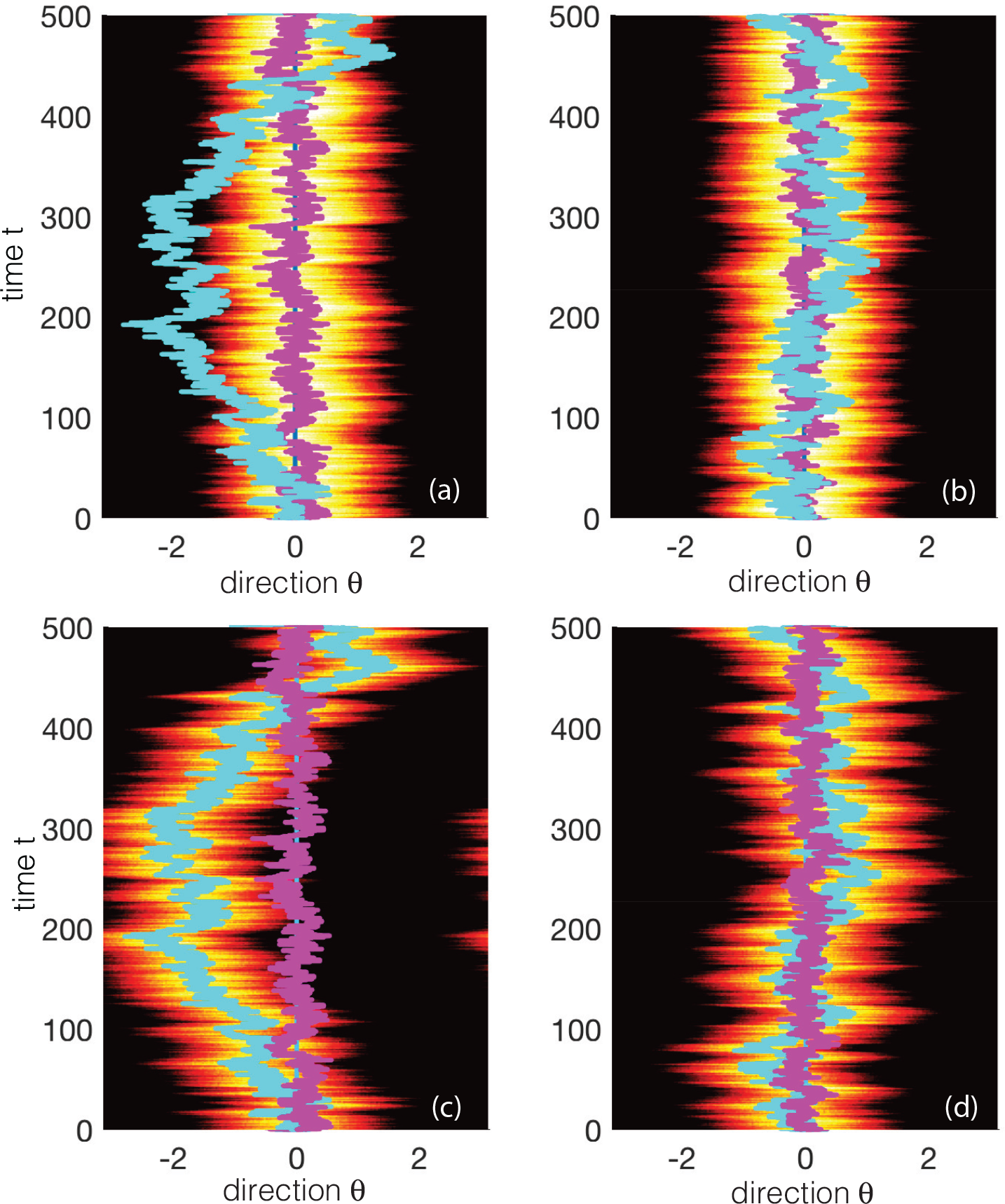
Same as Fig. 10 except that 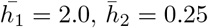 and 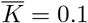 in (b,d).

In particular, for *θ* = *θ*′ we have

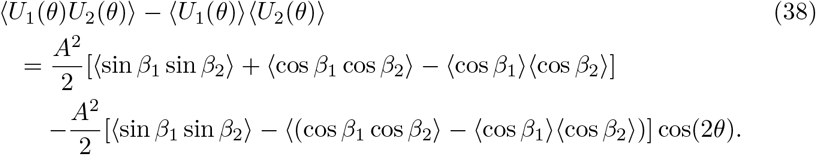

The resulting correlation tuning curve behaves in a similar fashion to the variance, see Fig. 9, where

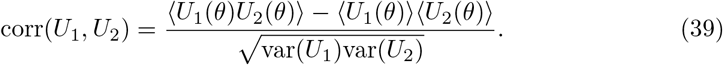

The above qualitative analysis can be confirmed by numerical simulations of the full neural field equations (1), as illustrated in Fig. 10 for a pair of identical ring networks. In Fig. 11, we show corresponding results for the case where network 2 receives a weaker stimulus than network 1 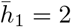 and 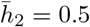. In the absence of interlaminar connections, the phase of network 2 fluctuates much more than the phase of network 1. When interlaminar connections are included, fluctuations are reduced, but network 2 still exhibits greater variability than network 1. This latter result is consistent with an experimental study of neural variability in V1 [74], which found that neural correlations were more prominent in superficial and deep layers of cortex, but close to zero in input layer 4. One suggested explanation for these differences is that layer 4 receives direct feedforward input from the LGN. Thus we could interpret network 1 in model A as being located in layer 4, whereas network 2 is located in a superficial layer, say.

### Effects of inter-network coupling (model B)

Our final example concerns a pair of coupled ring networks that represent horizontally connected center and surround hypercolumns, as shown in Fig. 2 (model B), with inter-network weight distribution (7). Again, for analytical tractability, we impose the symmetry conditions *A*_1_ = *A*_2_ = *A* and 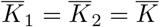. However, in contrast to model *A*, we take the contrasts to be the same,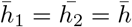, but allow the biases of the two inputs to differ, 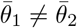. Equations (10) become, see **Materials and Methods**

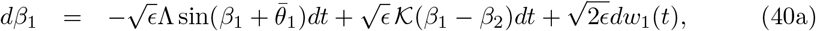

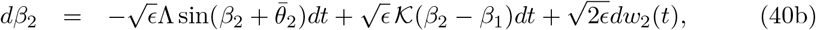

with *w*_*j*_ (*t*) given by equation (24) and

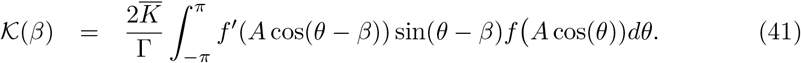

We can rewrite 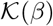 in the form

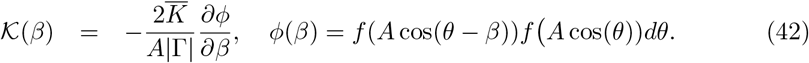

Note that *ϕ*(−*β*) = *ϕ*(*β*) and thus *ϕ*′(−*β*) = −*ϕ*′ (*β*). A sample plot of the potential *ϕ*(*β*) is shown in Fig. 12 together with an approximate curve fitting based on a von Mises distribution. For the given firing rate parameters *η* = 0.5 and *γ* = 4, the unperturbed bump amplitude is *A* ≈ 1.85.

As in the case of model A, we can rewrite equations (40) in the more compact form

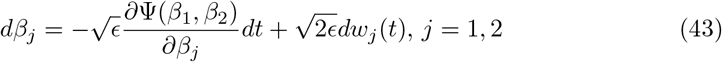

where Ψ is the potential function

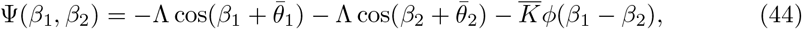

and we have absorbed the factor 2/(*A*|Γ|) into the constant 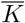. The corresponding two-dimensional forward Fokker-Planck equation is

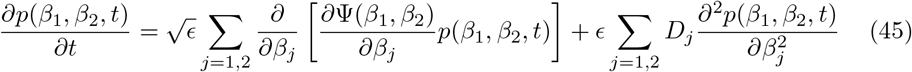

for *β*_*j*_ ∈ [−*π*, *π*] and periodic boundary conditions *p*(−*π*, *β*_2_, *t*) = *p*(*π*, *β*_2_, *t*), *p*(*β*_1_, −*π*, *t*) = *p*(*β*_1_, *π*, *t*). Following the analysis of model A, if *D*_1_ = *D*_2_ = *D* then the stationary density takes the form

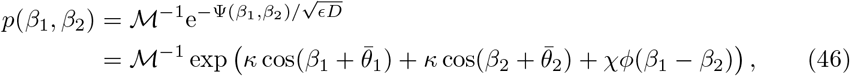

where

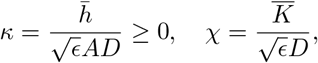

and 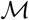 is a normalization factor.

Suppose that ring network 1 represents a hypercolumn driven by a center stimulus 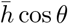 and network 2 represents a hypercolumn driven by a surround stimulus 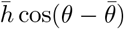, see Fig. 2. In Fig. 13 we plot how the normalized maximal variance of the center (at *θ* = *π*/2) varies with the directional bias 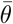 of the surround. We also show the baseline variance in the absence of horizontal connections (*χ* = 0). It can be seen that for an excitatory surround (*χ* > 0) the variance is suppressed in the center relative to baseline when the center and surround stimuli have similar biases 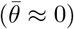 and is enhanced when they are sufficiently different 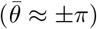. The converse holds for an inhibitory surround (*χ* < 0).

**Figure 12.**
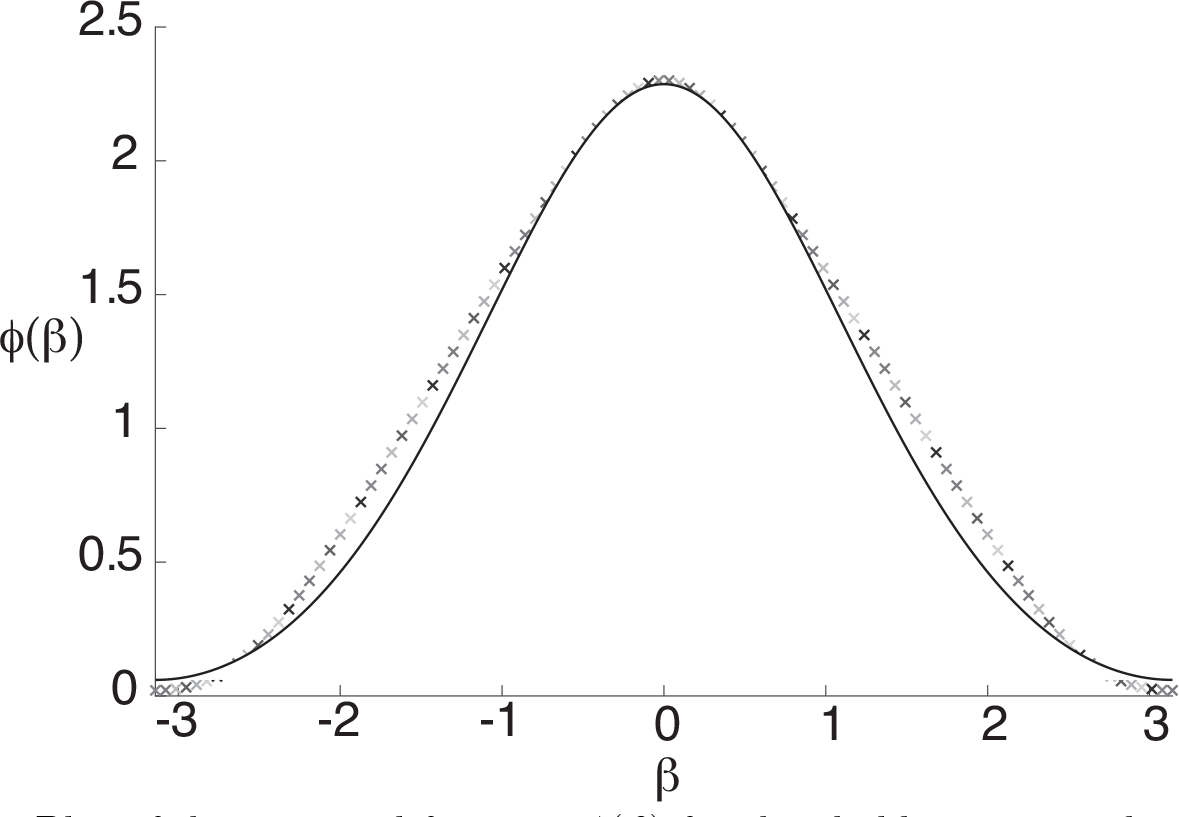
Plot of the potential function *ϕ*(*β*) for threshold *η* = 0.5 and gain *η* = 4. The solid curve is an approximation based on a fitted von Mises distribution *ϕ*(*β*) ≈ 12*M*(*β*; 0,0.6) − 0.9.

**Figure 13.**
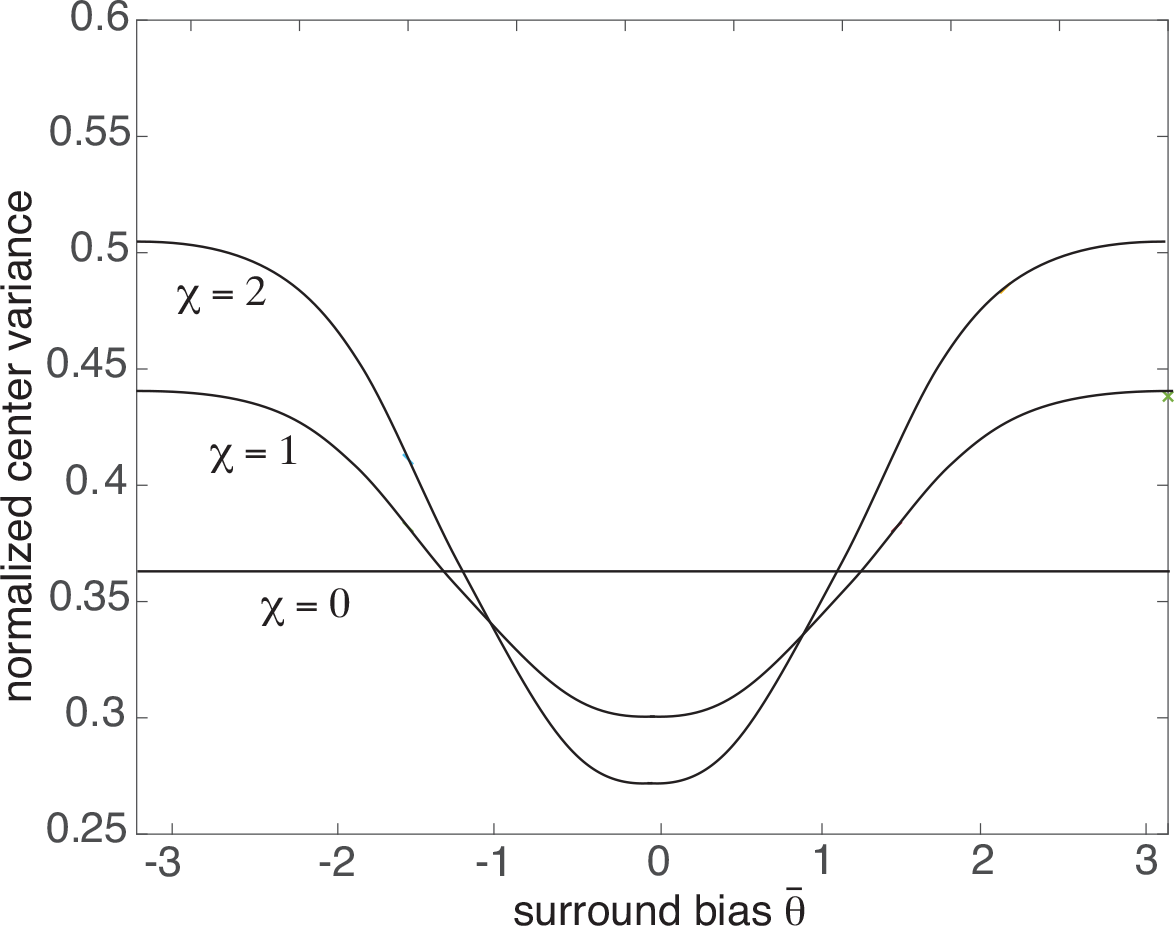
Coupled ring network (model B). Plot of normalized variance var(*U*_1_)/*A*^2^ of a center ring network as a function of the surround directional bias 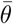 for various coupling parameters *χ*. Stimuli to center and surround are 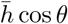 and 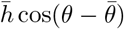, respectively, with 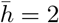.

## Discussion

In this paper we used stochastic neural field theory to analyze stimulus-dependent neural variability in ring attractor networks. In addition to providing a mathematical underpinning of previous experimental observations regarding the bimodal tuning of variability in directionally specific MT neurons, we also made a number of predictions regarding the effects of inter-network connections on noise suppression:

1. Excitatory vertical connections between cortical layers can suppress neural variability; different cortical layers can exhibit different degrees of variability according to the strength of afferents into the layers.
2. At low stimulus contrasts, surround stimuli tend to suppress (facilitate) neural variability in the center when the center and surround stimuli have similar (different) biases.
3. At high stimulus contrasts, surround stimuli tend to facilitate (suppress) neural variability in the center when the center and surround stimuli have similar (different) biases.

One of the main simplifications of our neural field model is that we do not explicitly model distinct excitatory and inhibitory populations. This is a common simplification of neural fields, in which the combined effects of excitation and inhibition are incorporated using, for example, Mexican hat functions [10, 30]. In the case of the ring network, the spontaneous formation of population orientation tuning curves or bumps is implemented using a cosine function, which represents short-range excitation and longer-range inhibition around the ring. We note, however, that the methods and results presented in this paper could be extended to the case of separate excitatory and inhibitory populations, as well as different classes of interneuron [9, 71].

One final comment is in order. Neural variability in experiments is typically specified in terms of the statistics of spike counts over some fixed time interval, and compared to an underlying inhomogeneous Poisson process. Often Fano factors greater than one are observed. In this paper, we work with stochastic firing rate models rather than spiking models, so that there is some implicit population averaging involved. In particular, we focus on the statistics of the variables *u*_*j*_(*x*, *t*), which represent the activity of local populations of cells rather than of individual neurons, with *f* (*u*_*j*_) the corresponding population firing rate [10]. This will allow us to develop an analytically tractable framework for investigating how neural variability depends on stimulus conditions within the attractor model paradigm. In order to fit a neural field model to single-neuron data, one could generate spike statistics by taking *f*(*u*_*j*_) to be the rate of an inhomogeneous Poisson process. Since *f* (*u*_*j*_) is itself stochastic, this would result in a doubly stochastic Poisson process, which is known to produce Fano factors greater than unity [24]. Moreover, the various phenomena identified in this paper regarding stimulus-dependent variability would carry over to a spiking model, at least qualitatively.

## Materials and methods

We present the details of the derivation of the stochastic phase equations (10).

### Stationary bumps in a single uncoupled ring

First, suppose that there are no external inputs, no inter-network coupling (*J*_12_ = *J*_21_ = 0), and no noise (*ϵ* = 0). Each network can then be described by a homogeneous ring model of the form

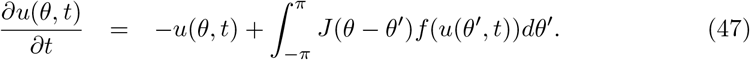

Let 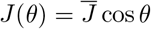 and consider the trial solution *u*(*θ*, *t*) = *U*(*θ*) with *U*(*θ*) an even, unimodal function of *θ* centered about *θ* = 0. This could represent a direction tuning curve in MT ((in the marginal regime) or a stationary bump encoding a spatial working memory. It follows that *U*(*θ*) satisfies the integral equation

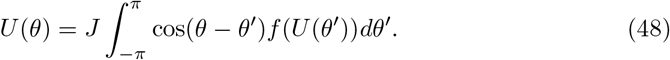

Substituting the cosine series expansion
 
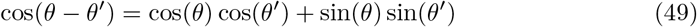

into the integral equation yields the even solution *U*(*θ*) = *A* cos *θ* with the amplitude *A* satisfying the self-consistency condition

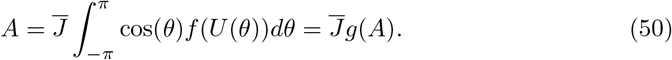

**Figure 14.**
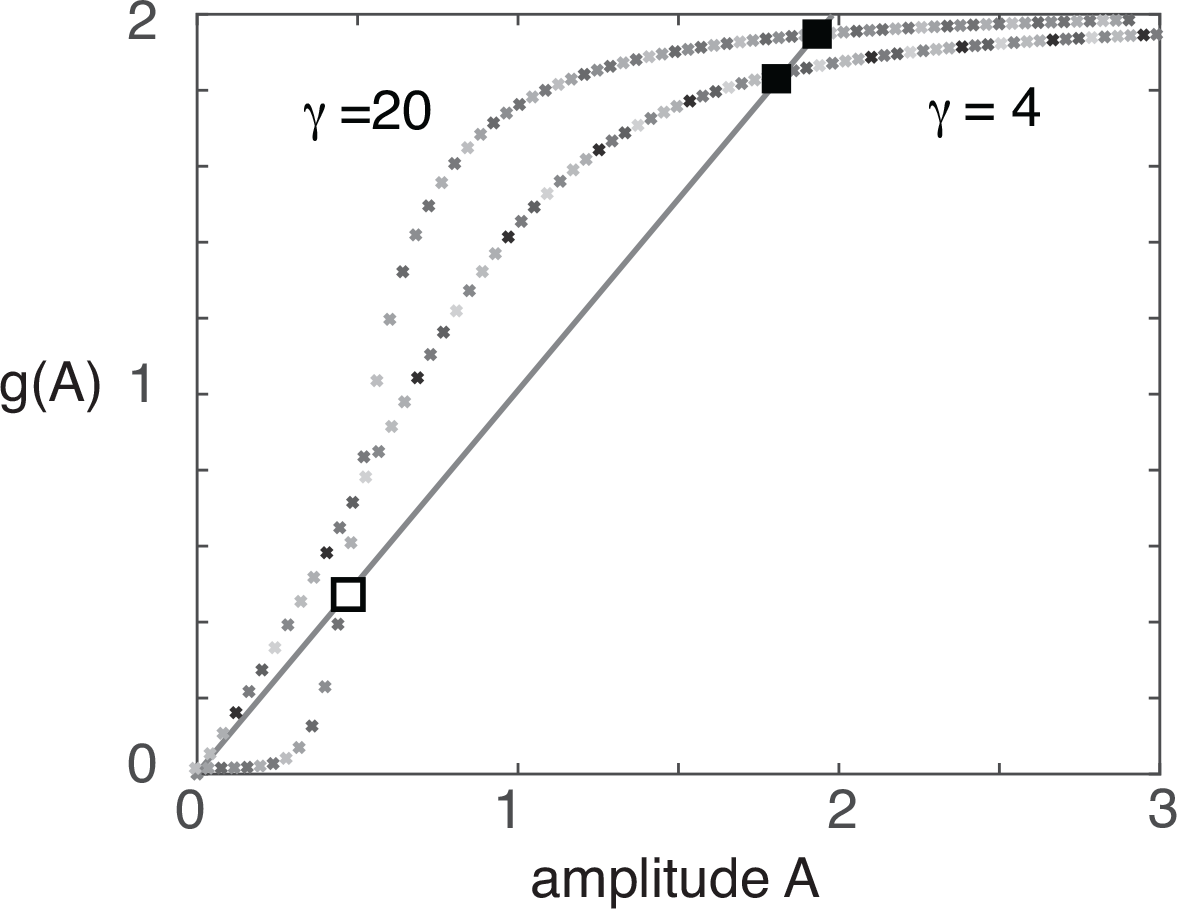
Graphical solution of the bump amplitude equation (50) for 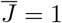 and *η* = 0.5. At intermediate gains (*γ* = 4) the zero solution is unstable and there exists a single stable bump. In the high gain limit (*γ* = 20) the zero solution is stable, and coexists with a small amplitude unstable bump and a large amplitude stable bump.

The amplitude equation (50) can be solved explicitly in the large gain limit *γ* → ∞, for which *f*(*u*) → *H*(*u* − *κ*), where *H* is the Heaviside function [46]. That is, 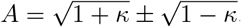, corresponding to a marginally stable large amplitude wide bump and an unstable small amplitude narrow bump, consistent with the original analysis of Amari [2]. On the other hand, at intermediate gains, there exists a single stable bump rather than an unstable/stable pair of bumps, see Fig. 14.

Linear stability of the stationary solution can be determined by considering weakly perturbed solutions of the form *u*(*θ*, *t*) = *U*(*θ*) + *ψ*(*θ*)e^*λt*^ for |*ψ*(*θ*)| ≪ 1. Substituting this expression into equation (47), Taylor expanding to first order in *ψ*, and imposing the stationary condition (48) yields the infinite-dimensional eigenvalue problem [9]

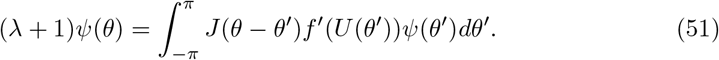

This can be reduced to a finite-dimensional eigenvalue problem by applying the expansion (49):

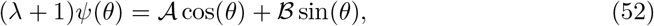

where

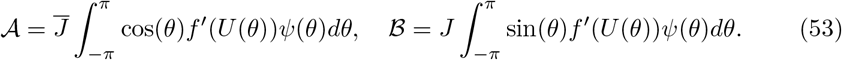

Substituting equation (52) into (53) then gives the matrix equation [46]

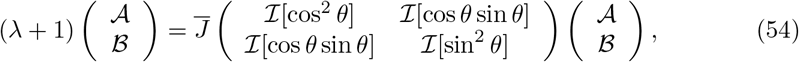

where for any periodic function *v*(*θ*)

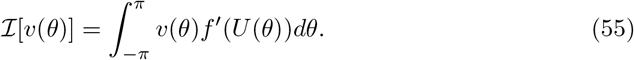

Integrating equation (50) by parts shows that for *A* ≠ 0

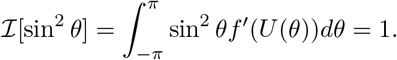

Hence, exploiting the fact that 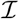 is a linear functional of *v*,

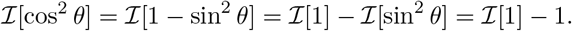

Finally, integration by parts establishes that

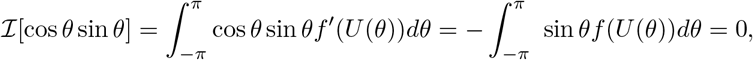

since *U*(*θ*) is even. Equation (54) now reduces to

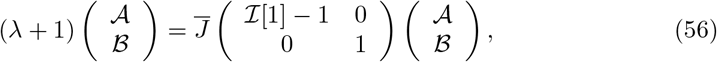

which yields the pair of solutions

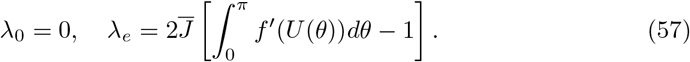

The zero eigenvalue is a consequence of the fact that the bump solution is marginally stable with respect to uniform shifts around the ring; the generator of such shifts is the odd function sin *θ*. The other eigenvalue *λ*_*e*_ is associated with the generator, cos *θ*, of expanding or contracting perturbations of the bump. Thus linear stability of the bump reduces to the condition *λ*_*e*_ < 0. This can be used to determine the stability of the pair of bump solutions in the high-gain limit [46].

A variety of previous studies have shown how breaking the underlying translation invariance of a homogeneous neural field by introducing a nonzero external input stabilizes wave and bump solutions to translating perturbations [31, 36, 37, 46, 81]. For the sake of illustration, suppose that 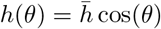 in the deterministic version of equation (1). This represents a weak *θ*-dependent input with a peak at *θ* = 0.

Extending the previous analysis, one finds a stationary bump solution 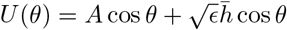, with *A* satisfying the implicit equation

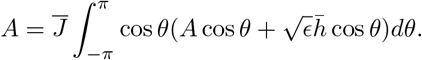

Again, this can be used to determine both the width and amplitude of the bump in the high-gain limit. Furthermore, the above analysis can be extended to establish that, for weak inputs, the bump is stable (rather than marginally stable) with respect to translational shifts [46].

### Perturbation analysis

The amplitude phase decompositions (*β*_*j*_, *v*_*j*_) defined by equation (9) are not unique, so additional mathematical constraints are needed in order to uniquely specify the decomposition, and this requires specifying the allowed class of functions of *v*_*j*_ (the appropriate Hilbert space). We will take take *v*_*j*_ ∈ *L*^2^(*S*^1^), that is, *v*_*j*_ (*θ*) is a periodic function with 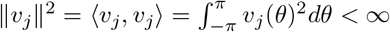. Substituting the decomposition into the stochastic neural field equation (1) and using Ito’s lemma gives [39]

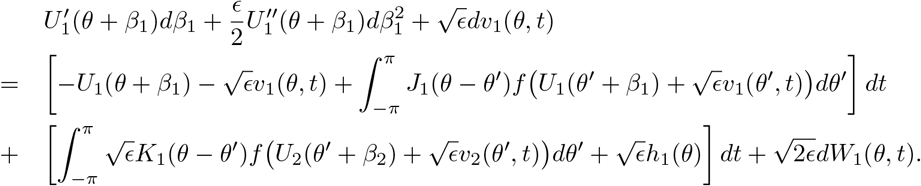

and

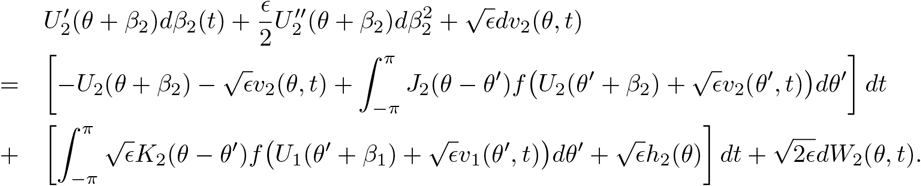

Introduce the series expansions 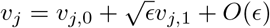, Taylor expanding the nonlinear function *F*, imposing the stationary solution (48), and dropping all *O*(*ϵ*) terms. This gives [10, 46], after dropping the zero index on *v*_*j*,0_,

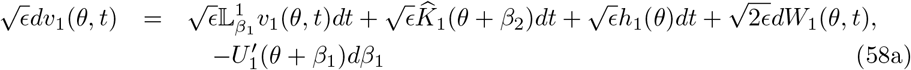

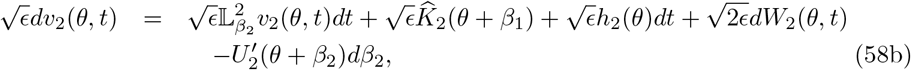

where 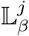 are the following linear operators

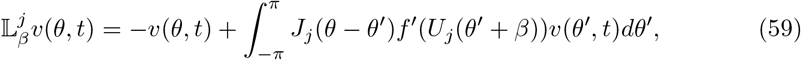

and

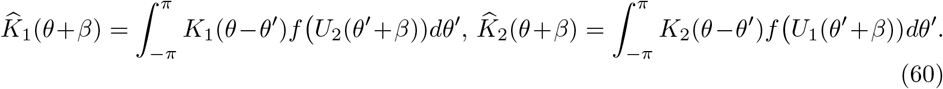

It can be shown that the operator 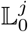 has a 1D null space spanned by 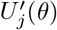. The fact that 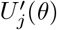 belongs to the null space follows immediately from differentiating equation (48) with respect to *θ*. Moreover, 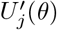 is the generator of uniform translations around the ring, so that the 1D null space reflects the marginal stability of the bump solution. (Marginal stability of the bump means that the linear operator 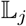 has a simple zero eigenvalue while the remainder of the discrete spectrum lies in the left-half complex plane. The spectrum is discrete since *S*^1^ is a compact domain.) This then implies a pair of solvability conditions for the existence of bounded solutions of equations (58a), namely, that *d ν*_*j*_ is orthogonal to all elements of the null space of the adjoint operator 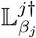. The corresponding adjoint operator is

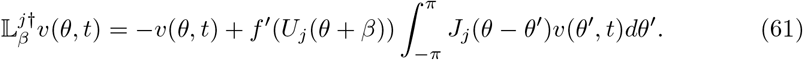

Let 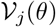 span the 1D adjoint null space of 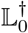. Now taking the inner product of both sides of equation (58a) with respect to 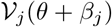 and using translational invariance then yields the following SDEs to leading order:

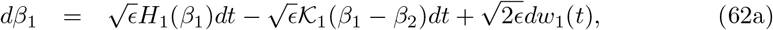

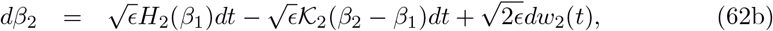

where

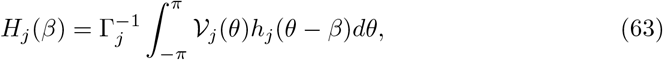

for *H*_*j*_(*β* + 2*π*) = *H*_*j*_(*β*),

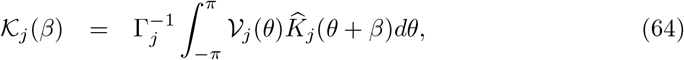

and

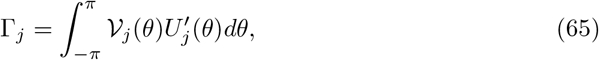

Here *w*_*j*_(*t*) are scalar independent Wiener processes,

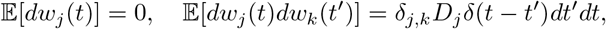

with

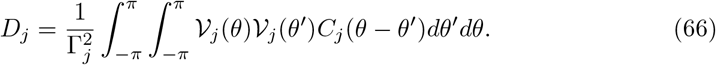

### Evaluation of functions *H*_*j*_ and 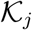

In order to determine the functions *H*_*j*_ and 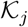 we need to obtain explicit expressions for the null vectors 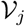. We will take 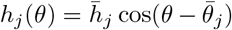. Applying the expansion (49) to the adjoint equation 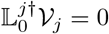 with 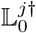 defined by equation (61), we can write [46]

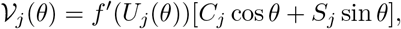

with

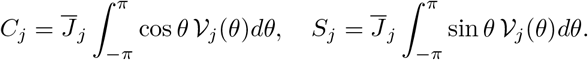

Substituting the expression for 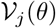 into the expressions for *C*_*j*_ and *S*_*j*_ then leads to a matrix equation of the form (56) with *λ* = 0. Since 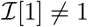, it follows that *C*_*j*_ = 0 so that, up to scalar multiplications,

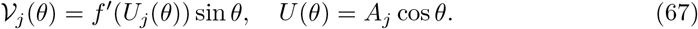

Now substituting 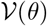 into equation (63), we have

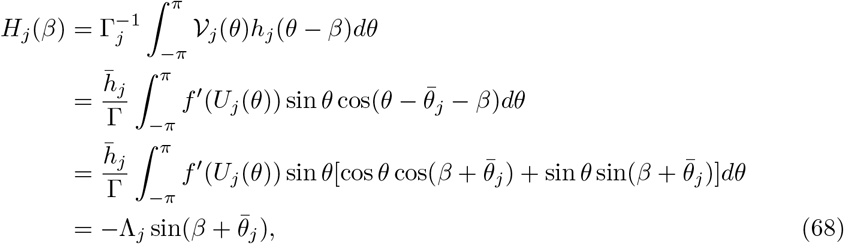

with

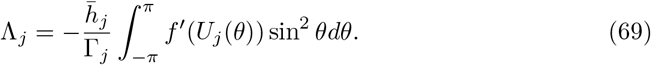

We have used the fact that *f*″(*U*_*j*_(*θ*)) is an even function of *θ*, so that the coefficient for 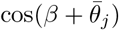 is zero. The constant Γ_*j*_ can be calculated from equation (65):

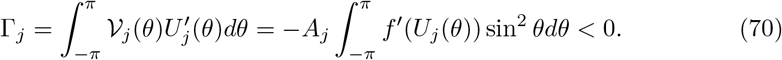

It follows that

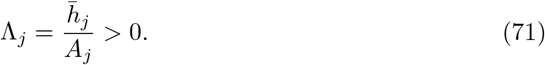

The calculation of 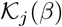 depends on whether we consider model A or model B, see Figs. 14 and 2. From equations (6), (60) and (64), we have for model A

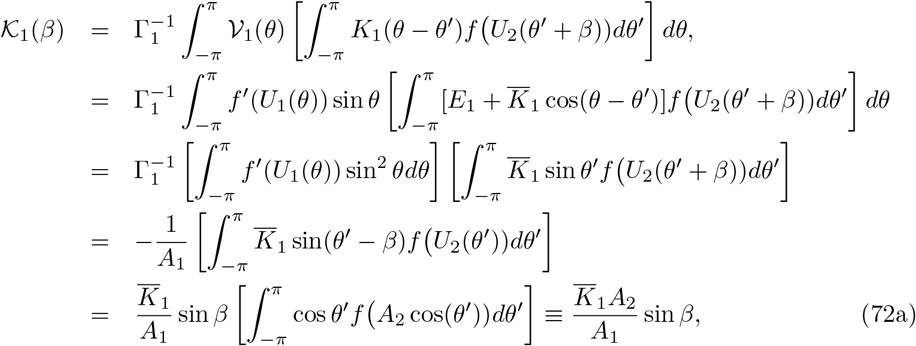

where we have used the stationary condition (8), and

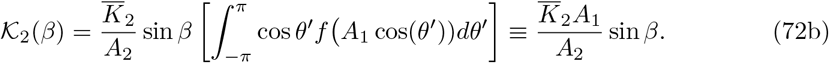

Similarly, from equations (7), (60) and (64), we have for model B

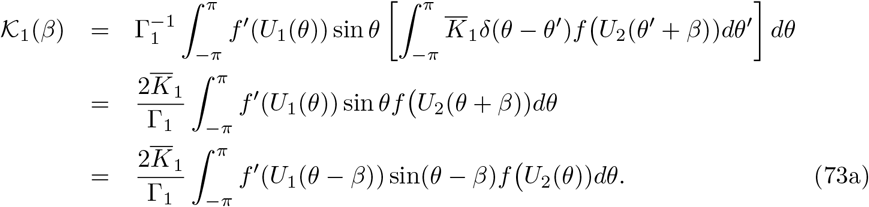

Similarly,

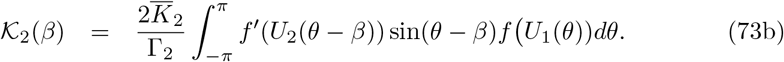

### Evaluation of diffusion coefficients

Finally, from equation (66), the diffusion coefficients *D*_*j*_ become

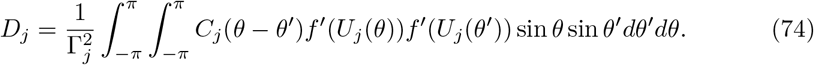

One finds that the diffusivities decreases as the spatial correlation lengths increase. For example, in the case of spatially homogeneous noise 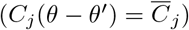, *D*_*j*_ = 0 since *f*′ (*U*_*j*_ (*θ*)) is even. On the other hand, for spatially uncorrelated noise 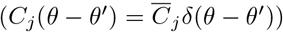, we have

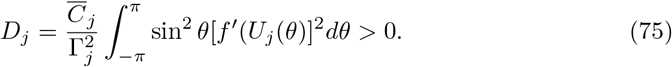

In **Results** we take 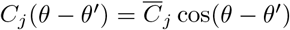 so that

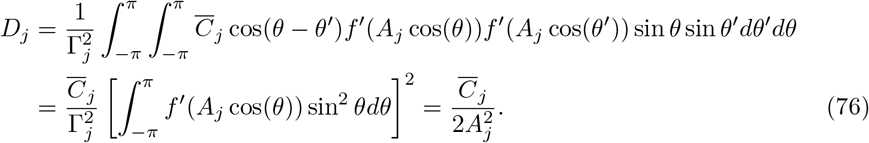

### Numerical methods

All numerical simulations were performed in Matlab. One dimensional numerical simulations were per-formed using a forward Euler method scheme in time and a trapezoidal rule for integration in *θ*. Time steps were taken to be Δ*t* = 0.001, and orientation steps Δ*θ* = 0.01*π*.

